# Profiling immunoglobulin repertoires across multiple human tissues by RNA Sequencing

**DOI:** 10.1101/089235

**Authors:** Serghei Mangul, Igor Mandric, Harry Taegyun Yang, Nicolas Strauli, Dennis Montoya, Jeremy Rotman, Will Van Der Wey, Jiem R. Ronas, Benjamin Statz, Douglas Yao, Alex Zelikovsky, Roberto Spreafico, Sagiv Shifman, Noah Zaitlen, Maura Rossetti, K. Mark Ansel, Eleazar Eskin

## Abstract

Assay-based approaches provide a detailed view of the adaptive immune system by profiling immunoglobulin (Ig) receptor repertoires. However, these methods carry a high cost and lack the scale of standard RNA sequencing (RNA-Seq). Here we report the development of ImReP, a novel computational method for rapid and accurate profiling of the immunoglobulin repertoire from regular RNA-Seq data. ImReP can also accurately assemble the complementary determining regions 3 (CDR3s), the most variable regions of Ig receptors. We applied our novel method to 8,555 samples across 53 tissues from 544 individuals in the Genotype-Tissue Expression (GTEx v6) project. ImReP is able to efficiently extract Ig-derived reads from RNA-Seq data. Using ImReP, we have created a systematic atlas of 3.6 million Ig sequences across a broad range of tissue types, most of which have not been studied for Ig receptor repertoires. We also compared the GTEx tissues to track the flow of Ig clonotypes across immune-related tissues, including secondary lymphoid organs and organs encompassing mucosal, exocrine, and endocrine sites, and we examined the compositional similarities of clonal populations between these tissues. The Atlas of Immunoglobulin Repertoires (The AIR), is freely available at https://github.com/smangul1/TheAIR/wiki, is one of the largest collection of CDR3 sequences and tissue types. We anticipate this recourse will enhance future immunology studies and advance the development of therapies for human diseases. ImReP is freely available at https://github.com/mandricigor/imrep/wiki

## Introduction

A key function of the adaptive immune system is to mount protective memory responses to a given antigen. B cells recognize their specific antigens through their surface antigen receptors (immunoglobulins, Ig), which are unique to each cell and its progeny. A typical Ig repertoire is composed of one immunoglobulin heavy chain (IGH) and two light chains, kappa, and lambda (IGK and IGL). Igs are diversified through somatic recombination, a process that randomly combines variable (V), diversity (D), and joining (J) gene segments, and inserts or deletes non-templated bases at the recombination junctions^1^ (Figure 1a). The resulting DNA sequences are then translated into antigen receptor proteins. This process allows for an astonishing diversity of the Ig repertoire (i.e., the collection of antigen receptors of a given individual), with >10^13^ theoretically possible distinct Ig receptors^1^. This diversity is key for the immune system to confer protection against a wide variety of potential pathogens ^2^. In addition, upon activation of a B cell, somatic hypermutation further diversifies Ig in their variable region. These changes are mostly single-base substitutions occurring at extremely high rates (10^−5^ to 10^−3^ mutations per base pair per generation)^3^. Isotype switching is another mechanism that contributes to B-cell functional diversity. Here, antigen specificity remains unchanged while the heavy chain VDJ regions join with different constant (C) regions, such as IgG, IgA, or IgE isotypes, and alter the immunological properties of a BCR. The pairing of heavy and light occurring in polyclonally activated B cells chains is another mechanism to increase the Ig diversity.

**Figure 1.**
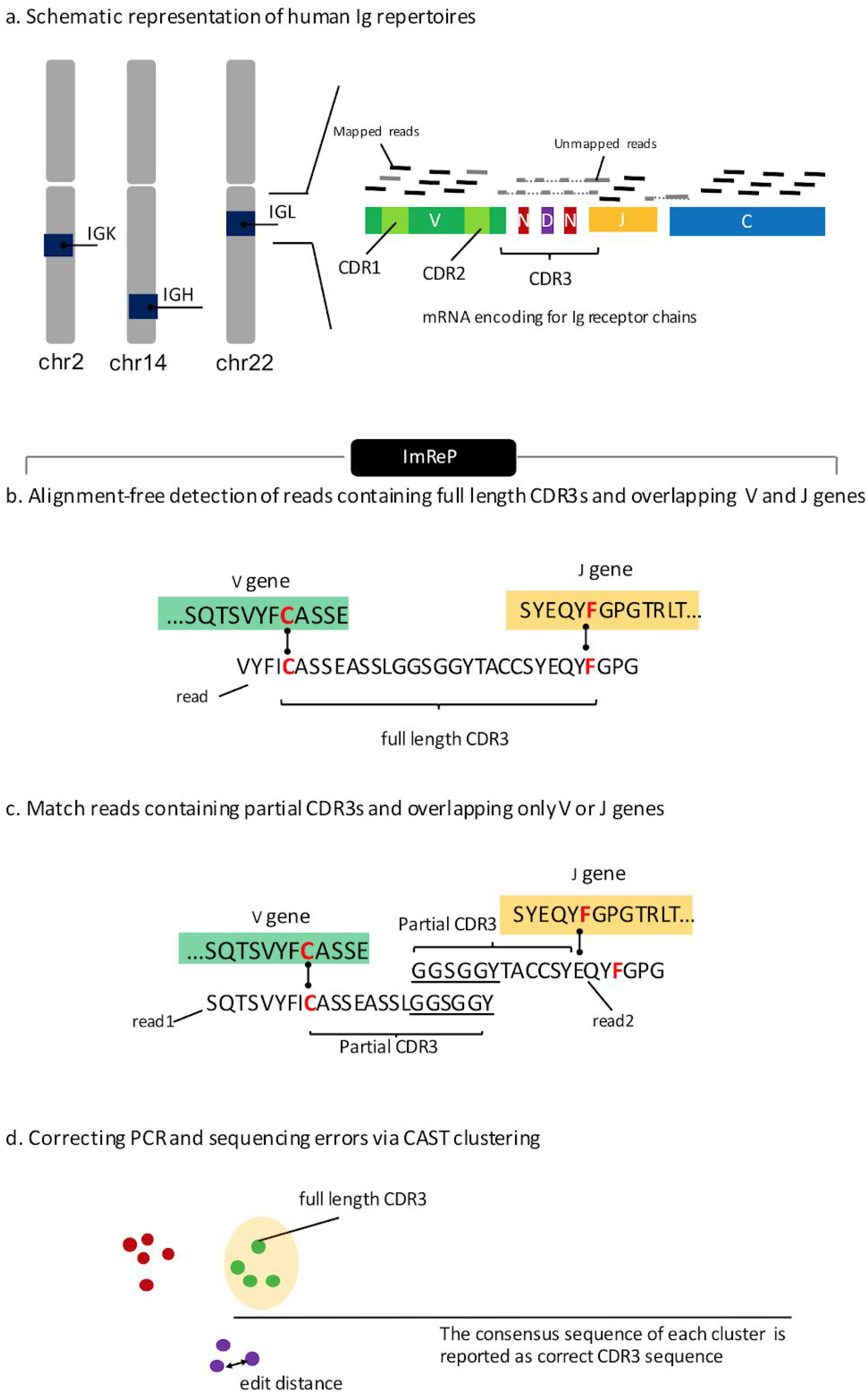
Overview of ImReP. **(a)** Schematic representation of human Ig receptor repertoire. Ig repertoire consists of three immunoglobulin loci (red color, Immunoglobulin heavy locus (IGH); Immunoglobulin kappa locus (IGK); Immunoglobulin lambda locus (IGL). Ig receptors contain multiple variable (V, green color), diversity (D, present only in IGH, violet color), joining (J, yellow color) and constant (C, blue color) gene segments. V(D)J gene segments are randomly jointed and non-templated bases (N, dark red color) are inserted at the recombination junctions. The resulting spliced Ig repertoire transcript incorporates the C segment and is translated into the antigen receptor proteins. RNA-Seq reads are derived from the rearranged immunoglobulin Ig loci. Reads entirely aligned to IG genes are inferred from mapped reads (black color). Reads with extensive somatic hypermutations and reads spanning the V(D)J recombination are inferred from the unmapped reads (grey color). Complementarity determining region 3 (CDR3) is the most variable region of the three CDR regions and is used to identify Ig receptor clonotypes—a group of clones with identical CDR3 amino acid sequences. **(b)** Alignment-free detection of reads containing full-length CDR3s and simultaneously overlapping V and J genes. Receptor derived reads spanning V(D)J recombinations are identified from unmapped reads and assembled into the CDR3 sequences. We first scan the amino acid sequences of the read to determines putative CDR3 sequences fully contained inside the read. The CDR3 sequence is a sequence starting with cysteine (C) and ending with (F) (IGK and IGL) or tryptophan (W) (for IGH). Reads with putative CDR3s are further examined to simultaneously overlap V and J gene segments. The alignment between the read and V and J genes is found by matching the prefix and suffix of the read to match the suffix of V and prefix of J genes, respectively. **(c)** Match reads containing partial CDR3s and overlapping only V or J genes. In case a read contains a partial CDR3 sequence and overlaps with only the V or J gene, we perform the second stage of ImReP. During this stage we match reads originated from the same CDR3 based on 15 nucleotides overlap. **(e)** Correcting PCR and sequencing errors via CAST clustering. We further correct PCR and sequencing errors in the assembled CDR3s. ImReP clusters assembled CDR3 into a set of clusters via CAST algorithm. The consensus sequence of each cluster is reported as correct CDR3 sequence.

High-throughput technologies enable unprecedented accuracy when profiling the Ig repertoires. Commonly used assay-based approaches provide a detailed view of the adaptive immune system with deep sequencing of amplified DNA or RNA from the variable region of the Ig locus (BCR-Seq)^4–6^. Those technologies are usually restricted to one chain, with the majority of studies focusing on the heavy chain of Ig repertoire. Recent studies^2^ successfully applied assay-based approaches to characterize the immune repertoire of the peripheral blood. However, little is known about the immunological repertoires of other human tissues, including barrier tissues like skin and mucosae. Studies involving assay-based protocols usually have small sample sizes, thus limiting analysis of *intra*-individual variation of immunological receptors across diverse human tissues.

RNA Sequencing (RNA-Seq) traditionally uses the reads mapped onto human genome references to study the transcriptional landscape of both single cells and entire cellular populations. In contrast to assay-based protocols that produce reads from the amplified variable region of Ig locus, RNA-Seq is able to capture the entire cellular population of the sample, including B cells. However, due to the repetitive nature of the Ig locus, as well as the extreme level of diversity in Ig transcripts, most mapping tools are ill-equipped to handle *Ig* sequences. RNA-Seq was successfully used for analysis of highly clonal leukemic repertoires with high relative quantities of *Ig* transcripts^5^. Despite this, Ig transcripts often occur in sufficient numbers within the transcriptome of many tissues to characterize their respective Ig repertoires^7^. A number of methods^8–10^ were designed to assemble Ig and T cell receptor repertoires and have been applied across various public RNA-Seq datasets. Existing methods that are capable of assembling Ig repertoire sequences produce results with low accuracy (f-score<.2), which are potentially biased and may lead to inaccurate conclusions (**Figure 2a**).

**Figure 2.**
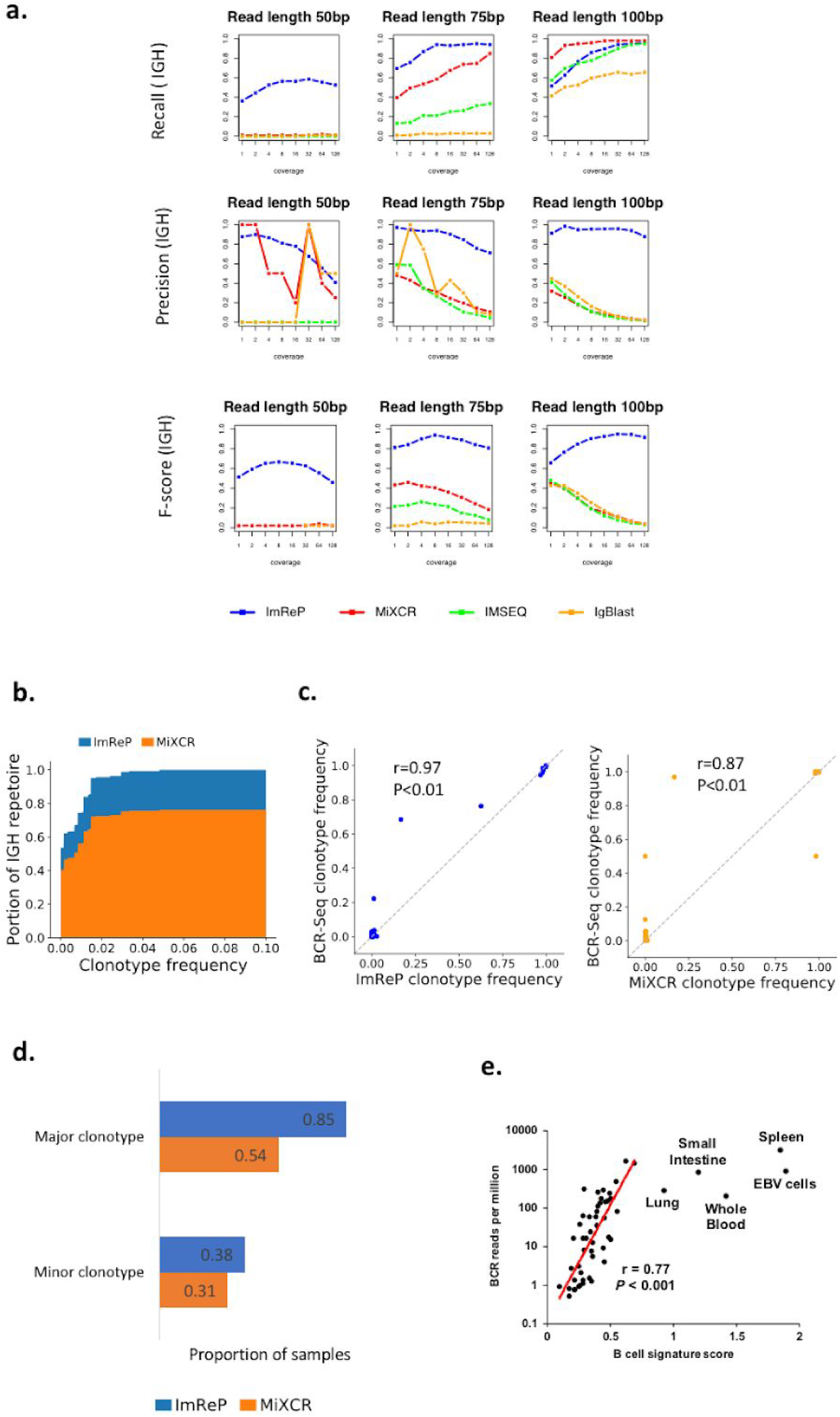
Evaluation of ImReP. **(a)** Evaluation of ImReP based on the number of assembled CDR3 sequences and comparison to the existing methods. Precision, recall and f-score rates for ImReP (blue), MiXCR (RNA-Seq mode) (red), IMSEQ (green), and IgBlast (orange) on simulated data for immunoglobulin heavy (IGH) transcripts are reported for various reads length (separate plots) and per transcript coverages (1,2,4,8,16,32,64,128) (x-axis). The recall was defined as TP/(TP+FN). Precision is defined as TP/(TP+FP). TP was defined as the number of correctly assembled CDR3 sequence (based on the exact match), FN - the number of true CDR3 sequence not assembled by the method, and FP - the number of incorrectly assembled CDR3 sequences. F-score was defined as the harmonic mean of precision and recall. Ig transcripts were simulated based on the random recombination of V and J gene segments (IMGT database) with non-template insertion at the recombination junction. **(b-d)** Concordance of targeted BCR-Seq and non-specific RNA-Seq performed on 13 tumor biopsies from Burkitt lymphoma. **(b)** Area chart shows the proportion of the total IGH repertoire captured by ImRep (blue) and MiXCR (RNA-Seq mode) (orange) depending on the minimum BCR-seq-confirmed clonotypes frequency considered. The *x-axis* corresponds to BCR-seq-confirmed clonotypes frequency Z. The *y*-axis corresponds to the fraction of assembled IGH repertoire with clonotype abundances greater than Z. The total repertoire was defined as the sum of the BCR-seq-confirmed clonotypes abundances. **(c)** Correlation of IGH clonotype frequencies estimated based on the BCR-Seq data (y-axis) and the RNA-seq data (x-axis) across all the samples. Only clonotypes assembled from RNA-Seq data are presented. (d) ImReP (blue) is able to detect major and minor clonotypes in a larger proportion of the samples compared to MiXCR (RNA-Seq mode) (orange). Major and minor clonotypes were defined based on BCR-Seq data as the clonotype with the largest frequency or smallest frequency, respectively. **(e)** Correspondence of ImReP-derived reads from Ig receptors to the relative abundance of B cells inferred from cell-specific gene expression profiles. Scatterplot of the number of all Ig-derived reads per 1 million RNA-Seq reads (y-axis) and B-cell signature score inferred by SaVant (x-axis).

In this study, we developed ImReP, a novel alignment-free computational method for rapid and accurate profiling of the Ig repertoire from regular RNA-Seq data. We applied it to 8,555 samples across 544 individuals from 53 tissues obtained from Genotype-Tissue Expression study (GTEx v6)^11^. The data was derived from 38 solid organ tissues, 11 brain subregions, whole blood, and three cell lines. ImReP is able to efficiently extract Ig-derived reads from the RNA-Seq data and accurately assemble the complementarity determining regions 3 (CDR3s). CDR3 are the most variable regions of the Ig receptors determining the antigen specificity. Using ImReP, we created a systematic atlas of Ig sequences across a broad range of tissue types, most of which were not previously studied for Ig repertoires. We also examined the compositional similarities of clonal populations between the tissues to track the flow of Ig clonotypes across immune-related tissues, including secondary lymphoid and organs that encompass mucosal, exocrine, and endocrine sites. Our proposed approach is not superior in comparison to targeted BCR-Seq; rather, it provides a useful tool for mining large-scale RNA-Seq datasets for the study of *Ig* receptor repertoires.

## Results

### Related work

A number of tools have been developed to reconstruct the *Ig* receptor repertoire. Repertoire analysis from RNA-Seq data typically starts with mapping the reads to the germline V, D, and J genes obtained from the International ImMunoGeneTics (IMGT) database^12^. There are three possible read mapping scenarios: (1) the read is entirely mapped to the V gene; (2) the read is entirely mapped to the J gene; (3) the read is partially mapped to the V and J genes simultaneously. Existing methods consider only reads from category (3). These methods use different underlying algorithms to map reads to germline genes. For example, MiXCR^8^ relies on an in-house alignment procedure, IgBlast^13^ utilizes BLAST with an optimized set of parameters, and IMSEQ^14^ uses in-house pairwise alignment between the read sequence and the germline V and J segment sequences.

Following the alignment, MiXCR performs overlapping of previously aligned reads into contigs. The resulting contigs are re-aligned to the V, D, and J genes to verify that the significant portion of non-template N insertions is covered. In contrast to MiXCR, which simultaneously aligns reads to both V, D, and J genes, IgBlast separately aligns the query read to the databases comprised of V, D, and J genes. IgBlast uses a specific sequence to separately align; first, the program finds the best V gene hit. Then, IgBlast masks the aligned read region and performs an alignment to the J gene database. (In the event of a heavy chain, IgBlast also queries the D gene database for the best hit.) The software checks that each component in the obtained V(D)J rearrangement originates from the same locus.

All methods use the definition of CDR3 to determine the boundaries of CDR3 sequence in each of the read. The last step in repertoire analysis is to correct the assembled clones for PCR and sequencing errors. In order to correct such errors, MiXCR and IMSEQ cluster the assembled clones and report a consensus sequence per cluster. IgBlast skips the error correction step and directly outputs inferred clones.

Most methods use alignment or assembly to infer CDR3s and align reads to V and J genes. In contrast, the ImReP procedure provides a match the read prefix and read suffix to the suffix of V and prefix of J genes, respectively, without using alignment. Avoiding alignment allows ImReP to significantly decrease running time and computational resources required to run the package. Average CPU time reported for ImReP is 44min; this runtime is substantially shorter than the 10 hours required for MiXCR. On average per sample, ImReP consumed 3G of CPU while MiXCR required 10G of CPU.

### ImReP: a two stage approach for immunoglobulin repertoires reconstruction

We applied ImReP to 0.6 trillion RNA-Seq reads (92 Tbp) from 8,555 samples to assemble CDR3 sequences IG receptors (**Table S2**). The RNA-Seq data was generated by the Genotype-Tissue Expression Consortium (GTEx v6). First, we mapped RNA-Seq reads to the human reference genome using a short-read aligner (performed by GTEx consortium11) (Figure 1). Next, we identify reads spanning the V(D)J junction of the IG receptors and assemble clonotypes (a group of clones with identical CDR3 amino acid sequences). We defined the CDR3 as the sequence of amino acids, starting with the cysteine on the left of the junction and ending with phenylalanine (for IGK, and IGL) or tryptophan (for IGH) on the right of the junction. Here ImReP used 0.02 trillion high quality reads that were successfully mapped to Ig genes or were unmapped reads that failed to map to the human reference genome (**Figures 1a and S1**).

ImReP is a two-stage alignment-free approach to assemble CDR3 sequences and detect corresponding V(D)J recombinations (**Figure 1b**). In the first stage, we prepare the candidate receptor reads from mapped and unmapped RNA-Seq reads (**Figure S1**). We merge partially mapped reads from BCR loci and unmapped reads into a set of candidate receptor reads, which serves as an input for ImReP. We scan the *amino acid* sequences of the read and determine the putative CDR3 as a substring of the read starting from *cysteine (C)* and ending with *phenylalanine (F)* (or *tryptophan (W)* for IGH). A read is decomposed in three parts: read prefix, CDR3, read suffix. The CDR3 sequence is a sequence starting with cysteine (C) and ending with (F) (IGK and IGL) or tryptophan (W) (for IGH). Reads with putative CDR3s are further examined to assess the overlap with V and J genes. Variable Ig receptor genes were imported from IMGT version: 3.1.17. We used C from the read (starting of CDR3) and C from the V gene (which usually occur as the three before the last codon in the V sequence) as an anchor to align the read prefix and V gene. Similarly, we used F (or W) from the read (end of CDR3) and F (or W) (which usually occur as the third codon in the J sequence). In the second stage, ImReP utilizes reads that contain a partial CDR3 sequence and overlap a single gene segment (V or J). Using the alignment-free procedure described above, we determined the alignment between the V or J gene and the read prefix or suffix, respectively. ImReP performs matching with a suffix tree technique. Matched reads with an overlap of at least 15 nucleotides are used to assemble full-length CDR3s. We further correct PCR and sequencing errors in the assembled CDR3s. ImReP clusters assembled CDR3 into a set of clusters via the CAST algorithm10. The clustering procedure is iteratively repeated until the average inverse edit distance (Levenshtein) inside each cluster is less than the user-defined threshold (ImReP default is .2). The consensus sequence of each cluster is reported as the correct CDR3 sequence. A detailed description of the methodology implemented with ImReP is provided in the Extended Experimental Procedures. ImReP is freely available at https://github.com/mandricigor/imrep/wiki. Currently, ImreP supports human and mouse *Ig* receptor repertoires.

### Feasibility of using RNA-Seq to study the *Ig* receptor repertoire

To validate the feasibility of using RNA-Seq to study the Ig receptor repertoire, we simulated RNA-Seq data as a mixture of transcriptomic reads and reads derived from IG transcripts (ratio between Ig-derived reads and transcriptomic reads was on average 1: 3600) (**Figure S2**). Ig transcripts are simulated based on random recombination of V, D and J gene segments (obtained from IMGT database^12^) with non-template insertion at the recombination junctions (**Figure S3**). We assessed the ability of ImReP to extract CDR3-derived reads from the RNA-Seq mixture by applying ImReP to a simulated RNA-Seq mixture. While our simulation approach may not completely summarize the various nuances and eccentricities of actual immune repertoires, it allows us to assess the accuracy of our tool. ImReP is able to identify 99% of CDR3-derived reads from the RNA-Seq mixture, suggesting it is a powerful tool for profiling RNA-Seq samples of immune-related tissues.

Next, we compared ImReP with other methods designed to assemble *Ig* receptor repertoires. We also investigated the sequencing depth and read length required to reliably assemble *Ig* sequences from RNA-Seq data. Our simulations suggest that both read length and sequencing depth have a major impact on precision-recall rates of CDR3 sequence assembly. ImReP is able to maintain an 80% precision rate for the majority of simulated scenarios. Average CDR3 coverage that is higher than 8 allows ImReP to archive a recall rate close to 90% for a read length above 75bp (**Figure 2a**). Increasing coverage has a positive effect on the number of assembled clonotypes achieved by ImReP.

We compared the performance of ImReP to MiXCR (RNA-Seq mode)^8,14^, IgBlast-based pipeline^15^, and IMSEQ^14^. Except for IMSEQ, these tools were developed to assemble the hypervariable sequences from Ig receptors directly from RNA-Seq data. Another tool, iSSAKE ^16^, is no longer supported and was not recommended for use. Unfortunately, we obtained empty output after running V’DJer^17^, and increasing coverage in the simulated data did not solve the problem. We exclude TRUST^9^ and TraCeR^10^, as those methods are solely designed for T cell receptors. We supplied each of those tools with the original RNA-Seq reads as raw or mapped reads, depending on the software developers’ recommendations. IMSEQ 14 cannot be applied directly to RNA-Seq reads because they were originally designed for targeted sequencing of *Ig* receptor loci. Thus, to independently assess and compare accuracy with ImReP, we only ran IMSEQ with the simulated reads derived from BCR transcripts (**Figure S2**). ImReP consistently outperformed existing methods in both recall and precision rates. The recall was defined as TP/(TP+FN). Precision was defined as TP/(TP+FP). We defined TP as the number of correctly assembled CDR3 sequence (based on the exact match), FN was defined as the number of true CDR3 sequence not assembled by the method, and FP was defined as the number of incorrectly assembled CDR3 sequences. On average, ImReP offers three-time superior accuracy (average f-score of ImRep was .78, for other methods average f-score was <.2). F-score was defined as a harmonic mean of precision and recall. Notably, ImReP was the only method with acceptable performance for 50bp read length, reconstructing with a higher precision rate significantly and more CDR3 clonotypes than other methods.

To further demonstrate the feasibility of applying non-specific RNA Sequencing to profile *Ig* receptor repertoires, we have used 18 tumor biopsies sequenced by BCR-Seq and RNA-Seq. Biopsies were acquired from the patients with histologically confirmed Burkitt lymphoma^18^. Per sample, 100 million paired-end RNA-Seq reads of length 50bp were available. We first mapped the reads onto the reference human genome and transcriptome and extracted unmapped reads, which were provided for ImReP to assemble IGH clonotypes. Based on the recommendation of the MiXCR, we have provided raw paired-end reads to the tool. BCR-Seq data was generated by Adaptive Biosystems (https://www.adaptivebiotech.com/) and was analyzed by Adaptive Biosystems’s Immune Analyzer package. One difficulty of using BCR-Seq as the gold standard to estimate the efficiency of the RNA-Seq method is that BCR-Seq captures DNA clonotypes, while RNA-Seq only captures the expressed clonotypes. To account for the possible discrepancies, we have first mapped RNA-Seq reads onto the major clonotypes with relative frequency at least 90% detected by BCR-Seq. In five of out 18 BCR-Seq samples, no RNA-Seq reads were mapped to BCR-seq-confirmed major clonotypes. Those samples were excluded from the analysis. In the remaining samples, we have considered the set of CDR3s obtained by BCR-Seq as the total IGH repertoire.

We investigated which portion of the total immune repertoire RNA-Seq is capable of capturing. Using RNA-Seq ImReP was able to capture on average 53.3% of the IGH repertoire, estimated as the sum of the detected BCR-seq-confirmed clonotypes. MiXCR was able to capture 40.1% respectively (**Figure 2b**). Overall, ImRep is capable to detect BCR-seq-confirmed clonotypes with a relative frequency exceeding 90% in all of the cases, while MiXCR in 83.3% of cases, respectively. When the frequency of the major clonotype drops below 10%, ImreP was able to detect the major clonotype in 60% of the cases, while MiXCR only in 20% of the cases, respectively. Remarkably both methods were able to detect major clonotype with the frequency below 1% in one of the samples (**Table S1**). We have also investigated the ability of both methods to detect BCR-seq-confirmed minor clonotypes. The average frequency of the minor clonotypes across all samples was 0.37%. ImreP was able to detect minor clonotype in 38% of the samples (**Figure 2d).** Despite the ability of ImreP and MiXCR to capture the majority of BCR-seq-confirmed repertoire, both methods often missed the rare clonotypes due to the limited number of BCR-derived reads in RNA-Seq data. ImReP was able to detect 50% of all BCR-Seq-confirmed clonotypes with the relative frequency higher than 0.24%. MiXCR was able to detect 50% of all BCR-Seq-confirmed clonotypes with the relative frequency higher than 0.29% (**Figure S4**). Both methods were able to accurately estimate the relative frequencies of assembled clonotypes (ImRep : r=0.97, p-value=10^-40^; MiXCR r=0.87, p-value=10^-15^) (**Figure 2c**). Scripts and commands utilized to process the data and run all tools used in this study are provided in the Extended Experimental Procedures and are available online at https://github.com/smangul1/imrep.GTEx/wiki.

We further validated the ability of ImReP to accurately infer the proportion of immune cells in the sampled tissue. We hypothesized that the fraction of B cells in the sample will be proportional with the fraction of receptor-derived reads in our RNA-Seq data. We used a transcriptome-based computational method, SaVant^19^, which uses cell-specific gene signatures (independent of *Ig* transcripts) to infer the relative abundance of B cells within each tissue sample (**Table S5**). The B cell signature used by SaVant are derived from CD19+ cells and might not represent every B cell subset^20^. However, CD19+ cells likely represent the largest populations of B cell subsets and many of the CD19 negative B cell subsets may still have a similar gene signature to the CD19 signatures. We found that B cell signatures inferred by SaVant showed the positive correlation with the size of IGH repertoire(r = 0.77, *P* < 0.001) (**Figure 2e**). An exception to this correlation was for tissues that contain the highest density of B cells: spleen, whole blood, small intestine (terminal ileum), lung, and EBV-transformed lymphocytes (LCLs).

### Characterizing the adaptive immune repertoire across 53 GTEx tissues

ImReP identified over 26 million reads overlapping 3.6 million distinct CDR3 sequences that originate from diverse human tissues. The majority of assembled CDR3 sequences derived from immunoglobulin heavy chain (IGH) (1.7 million), 0.9 million were derived from the immunoglobulin kappa chain (IGK), and 1.0 million from the immunoglobulin lambda chain (IGL). The vast majority of all assembled CDR3s had a low frequency in the data. 98% of CDR3 sequences had a count of less than 10 reads, and the median CDR3 sequence count was 1.4. CDR3 sequences derived from IGK were the most abundant across all tissues, accounting on average for 54% of the entire B-cell population (**Figure S5**).

We compared the length and amino acid composition^21^ of the assembled CDR3 sequences of *Ig* receptor chains (**Figure S6**). Consistent with previous studies, we observed that immunoglobulin light chains have notably shorter and less variable CDR3 lengths compared to heavy chains ^22^. The tissue type appears to have no effect on the length distribution of CDR3 sequences (**Figure S7**). In line with other studies ^22,23^, both light chains exhibited a reduced amount of sequencing diversity (**Figure S6**).

We observed per sample an average of 1331 distinct Ig clonotypes. We normalized the number of distinct clonotypes by the total number of raw RNA-Seq reads, which we call number of clonotypes per one million raw RNA-Seq reads (CPM). As the number of distinct clonotypes does not increase linearly with the sequencing depth, CPM metric should not be used in studies comparing clonotype diversity across various phenotypes. Instead, CPM is intended to be an informative measure of clonal diversity adjusted for sequencing depth. One technique allowing to properly adjust for sequencing depth per sample is to subsample reads, to have identical number of reads per sample.

We used per sample alpha diversity (Shannon entropy) to incorporate the total number of distinct clonotypes and their relative frequencies into a single diversity metric. Among all tissues, spleen has the largest B-cell population, with a median of 1301 Ig-derived reads per one million RNA-Seq reads. It also has the most diverse population of B cells with median per sample alpha diversity of 7.6 corresponding to 1025 CPM (**Figure 3** and **Table S2**). Organs that possess mucosal, exocrine, and endocrine sites (n=24) harbor a rich clonotype population with a median of 87 CPM per sample. Minor salivary glands have the highest IG diversity in the group (alpha=7.1) and surpass the diversity of the terminal Ileum containing Peyer’s Patches, which are secondary lymphoid organs (**Table S2**).

**Figure 3.**
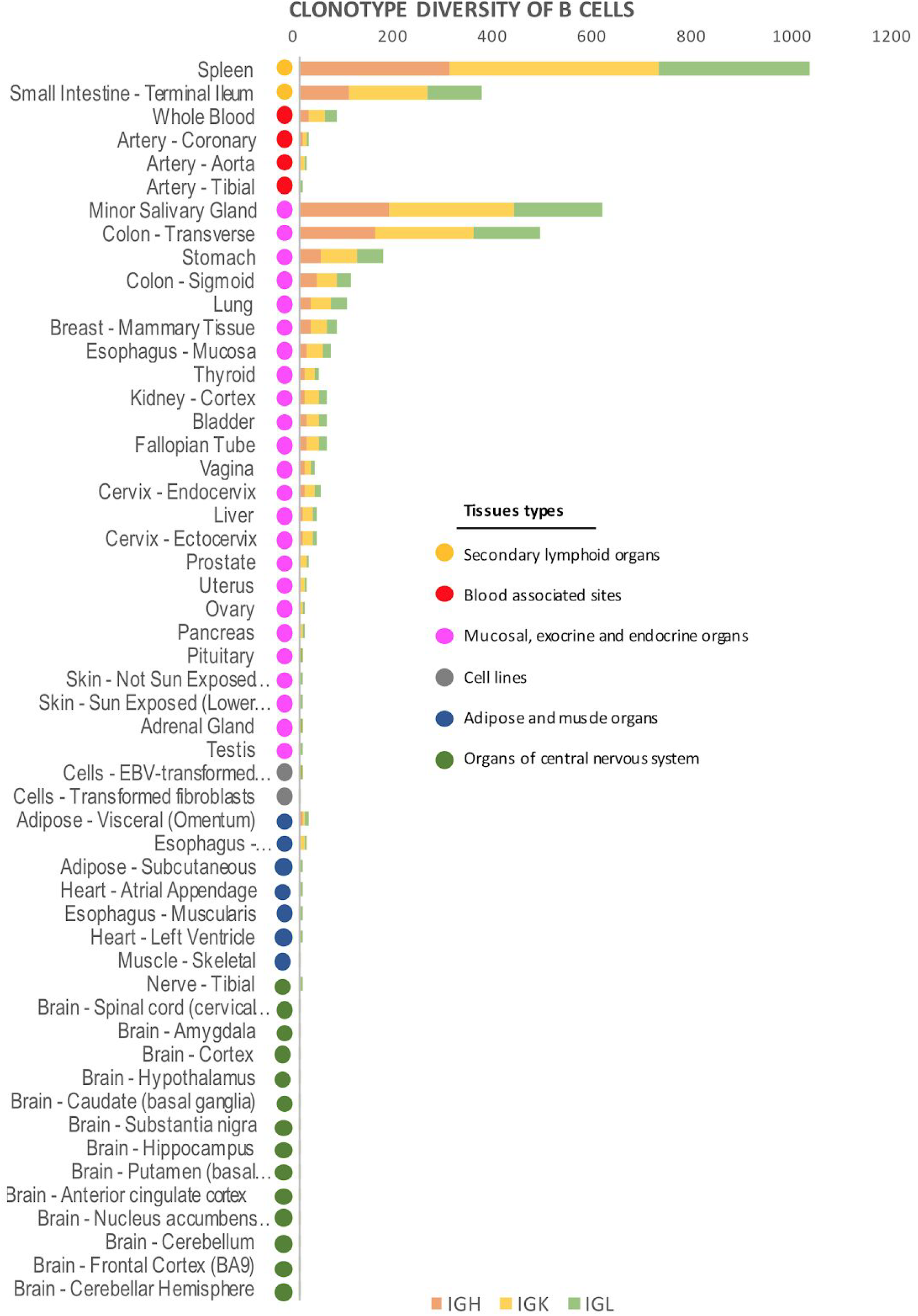
Adaptive immune repertoires across multiple human tissues. Adaptive immune repertoires of 8,555 samples across 544 individuals from 53 body sites obtained from Genotype-Tissue Expression study (GTEx v6). We group the tissues by their relationship to the immune system. The first group includes the lymphoid tissues (n=2, red colors). The second group includes blood associated sites including whole blood and blood vessel (n=4, red color). The third group are the organs that encompass mucosal, exocrine and endocrine organs (n=21, lavender color). The fourth group are cell lines (n=3, grey color). The fifth group are adipose or muscle tissues and the gastroesophageal junction (n=7, blue color). The sixth group are organs from the central nervous system (n=14, green color). Histogram reports clonotypic richness of B cells, calculated as the number of distinct amino acid sequences of CDR3 per one million RNA-Seq reads (CPM). The median number of distinct amino acid sequences of CDR3 are presented individually for immunoglobulin heavy chain (IGH), immunoglobulin kappa chain (IGK), immunoglobulin lambda chain (IGL).

Tissues not related to the immune system, including adipose, muscle, and the organs from the central nervous system, contained a median of 6 CPM per sample, which are most likely due to the blood content of the tissues24. The highest number of distinct CDR3 sequences among non-lymphoid organs was present in the omentum, a membranous double layer of adipose tissue containing fat-*associated lymphoid* clusters. As expected25, Epstein–Barr virus (EBV)-transformed lymphocytes (LCL) harbored a large homogeneous population of *Ig* clonotypes (**Table S2** and **Figure S8**). The number of reported clonotypes was normalized by the proportion of B cells within each tissue sample (**Table S3**). We have used SaVant to infer the relative abundance of B cells within each tissue sample based on cell-specific gene signatures (independent of *Ig* transcripts).

### Individual- and tissue-specific *Ig* clonotypes

Amino acid sequences of clonotypes exhibited extreme inter-individual dissimilarity, with 88% of clonotypes unique to a single individual (private) (**Figure 4a**). The remaining ~400,000 clonotypes were shared by at least two individuals (public). A small fraction of B cells in many tissues limits our ability to capture the entire Ig repertoire and thus resulting in classifying some public clonotypes as private. The number of individuals sharing clonotypes varied across Ig chains, with immunoglobulin light chains having the highest number of public clonotypes. Twenty-five percent of all IGK clonotypes were public, and the number of individuals sharing the IGK clonotype sequences can be as high as 471 (**Figure 4b**). The limited capacity of RNA-Seq to cover low abundant clonotypes may misclassify public clonotypes as private. Consistent with the previous studies^9,26^, we observe public clonotypes to be significantly shorter in length than the private ones (p-value<2×10^-16^). For example, IGH chain public clonotypes had an average length of 13 amino acids, and private clonotypes had an average length of 16. We also examined whether the public clonotypes were more often shared across tissues within an individual. Only 14% of the ~240,000 clonotypes shared across tissues were public. The majority of clonotypes were individual- and tissue-specific (**Figure 4c**). The full list of public clonotypes is distributed with the Atlas of Immunoglobulin Repertoires (TheAIR) that is publically available at https://github.com/smangul1/TheAIR/wiki.

**Figure 4.**
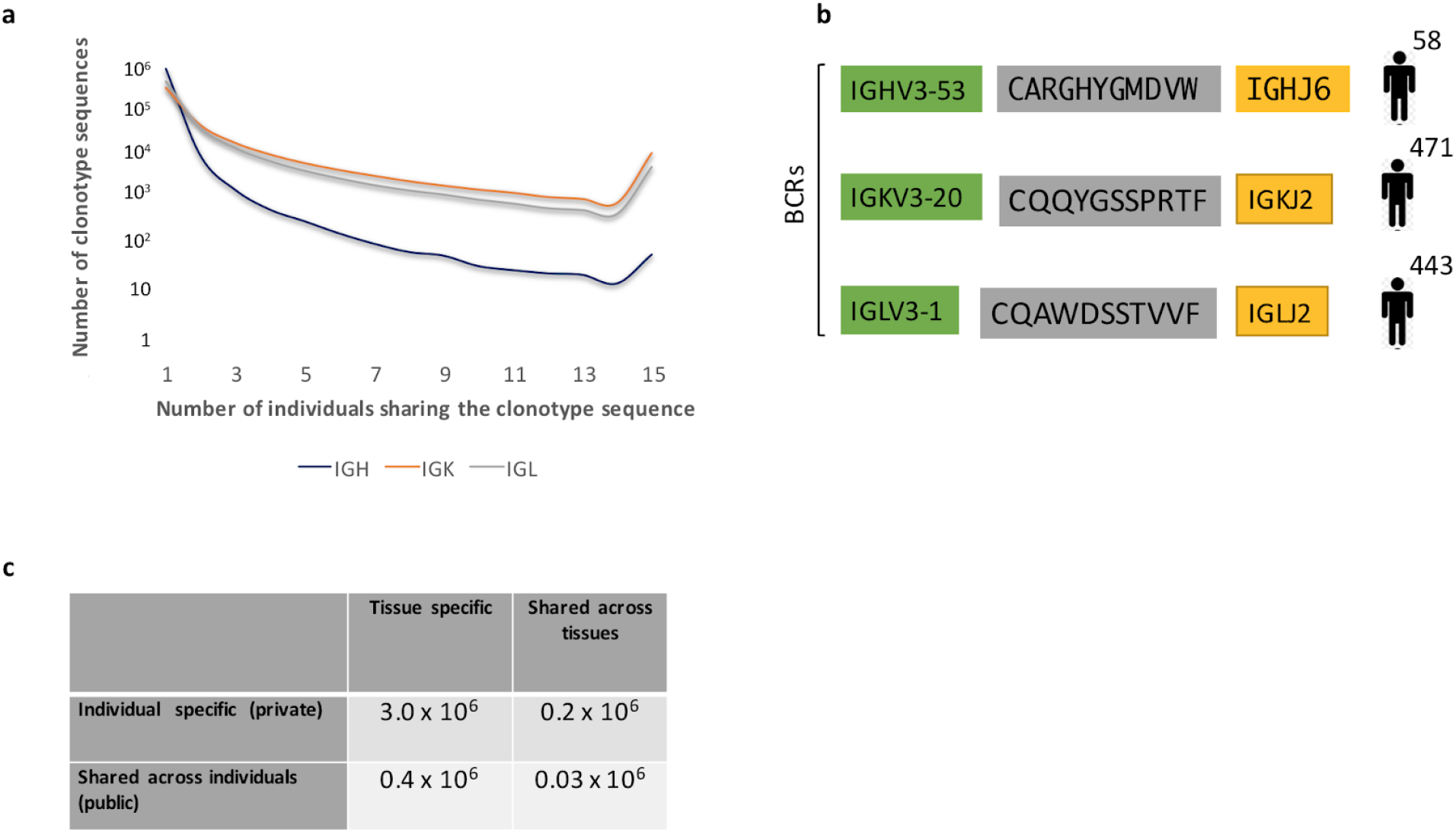
Private and public Ig clonotypes. (a) Distribution of frequencies of private (n=1) and public (n>1) clonotypes across 544 individuals. We collect clonotypes from all tissues of the same individual into a single set corresponding to that individual. (b) The most public clonotypes (shared across the maximum number of individuals) and corresponding VJ recombination are presented for IGH, IGK, IGL. (c) Clonotypes sequences are classified into public clonotypes (shared across individuals), private (individual-specific), tissue-specific, and clonotypes shared across multiple tissues. The number of clonotypes falling into each pair of categories is reported across Ig receptor chains.

### The flow of *Ig* clonotypes across human GTEx tissues

A large number of individuals available through the study allow us to establish a pairwise relationship between the tissues and track the flow of Ig clonotypes across human tissues. We observed a significant increase in the number of CDR3 sequences shared across pairs of tissues from the same individuals. Further, we observed this pattern consistently for all chains of Ig receptors (p-value<2×10^−16^) (**Figure 5a** and **Table S4**). We observe a different amount of shared CDR3 sequences across different types of Ig chains with an increase in immunoglobulin light chains compared to *Ig* heavy chain. On average, we observe 7.0 CDR3 sequences to be shared across a pair of tissues from the same individuals. Pairs of tissues from different individuals share on average 3.6 CDR3 sequences (**Figure 5a and Table S4**).

**Figure 5.**
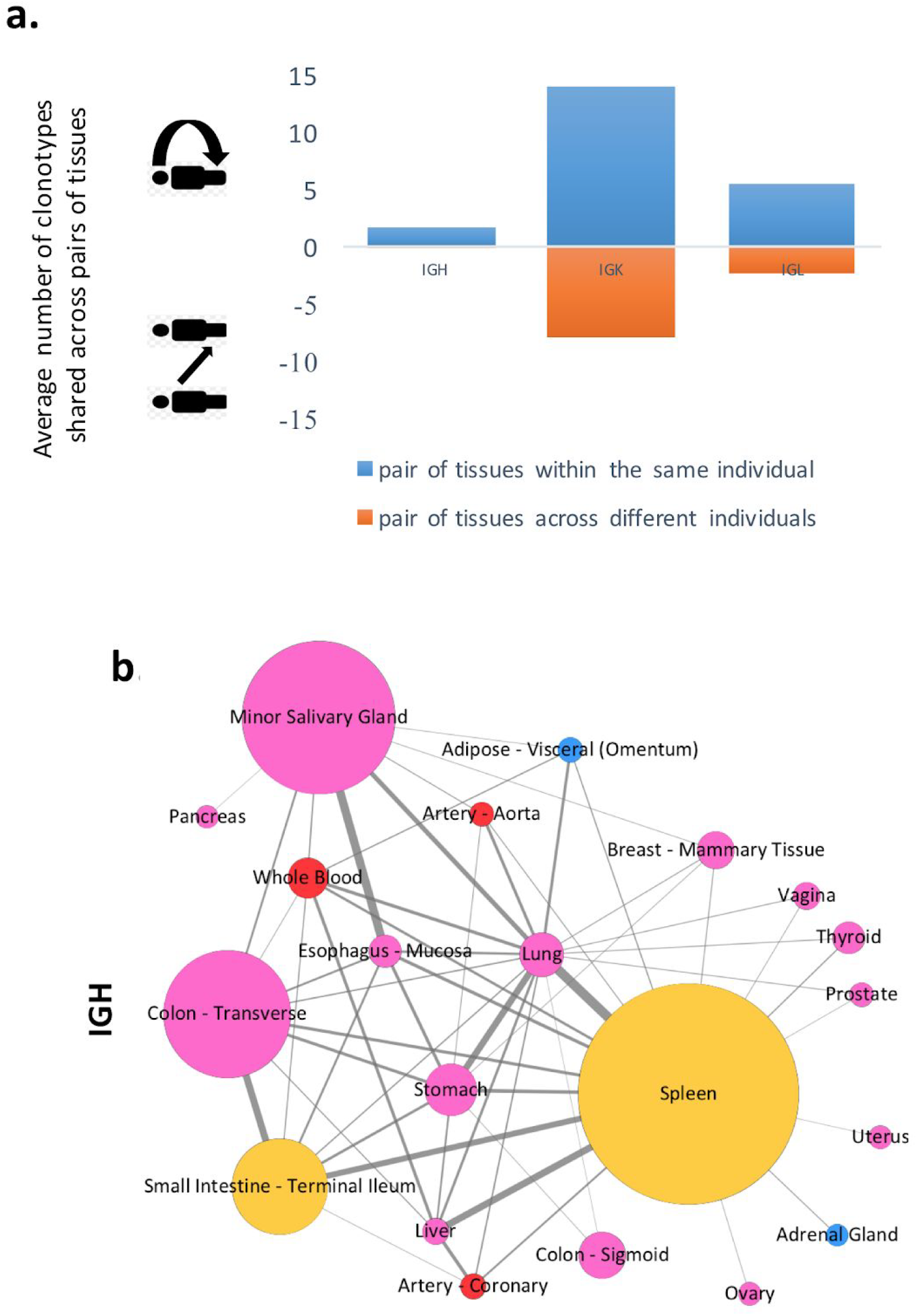
The flow of *Ig* clonotypes across diverse human tissues. Results are based on pairs of tissues with at least 10 individuals. **(a)** The number of *Ig* clonotype sequences shared across pairs of tissues from the same individuals (blue color) and from different individuals (orange color) is presented. **(b)-(c)** Flow of clonotypes across diverse human tissues is presented as a network. Each node is a tissue with the size proportional to a median number of clonotypes of the tissue. The color of the node corresponds to a type of the tissue type: lymphoid tissues (yellow colors), blood associated sites (red color), organs that encompass mucosal, exocrine and endocrine organs (lavender color). Compositional similarities between the tissues in terms of gain or loss of CDR3 sequences are measured across valid pairs of tissues using beta diversity (Sørensen–Dice similarity index). Edges are weighted according to the beta diversity. **(b)** The flow of *IGH* clonotypes across diverse human tissues is presented as a network. Edges with beta diversity >.001 are presented.

To establish the flow of *Ig* clonotypes across various tissues, we compared clonotype populations between and within the same individuals. We limited this analysis to pairs of tissues for which we had at least 10 individuals (870 pairs of tissues out of 1378 possible pairs). We used beta diversity (Sørensen–Dice similarity index) to measure compositional similarities between the tissues in terms of gain or loss of CDR3 sequences (**Figure 5b**). For the majority of the 870 available tissue pairs, we observe no *IGH* sequences in common, which corresponds to beta diversity of 0.0.

We have examined the flow of *IGH* clonotypes across tissues and presented it as a network (Figure 5b). Among 870 available tissue pairs, we have identified 56 tissue pairs with beta diversity above .001. The spleen was the most highly connected tissue, with 17 connections, followed by lung, with 16 connections. Clonotypes represents one connected component, meaning that every two nodes are connected directly or via other nodes. Clonotype populations of spleen and lung are the most similar (0.02 beta diversity), other pairs include minor salivary gland and esophagus mucosa, terminal ileum (small intestine) and transverse colon. We observe above 200 pairs of tissues with beta diversity above .001 for immunoglobulin light chains (**Figure S9-S10**). The most similar tissue pairs for *IGK* chain were spleen and transverse colon (0.15 beta diversity).

### ImReP identifies tissue samples with lymphocyte infiltration

Histological images of tissue cross-sections and pathologists’ notes have been used to validate the ImReP’s ability to detect the samples with a high lymphocyte content, which often corresponds to a disease state. We examined the IGH clonotype populations from thyroid tissue across individuals. The median number of inferred distinct CDR3 sequences per sample was 20, though 14.5% of the samples had more than 500 distinct CDR3 sequences. We observed the highest number of CDR3 sequences among all the thyroid samples in an individual with late-stage Hashimoto’s thyroiditis, an autoimmune disease characterized by lymphocyte infiltration and T-cell mediated cytotoxicity. According to pathologists’ notes, Hashimoto’s disease was present in 11.2% of thyroid samples, with varying degrees of severity. First, we used pathologists’ notes to annotate samples as healthy (n=183) or bearing Hashimoto’s disease (n=23), and then we compared the adaptive repertoire diversity between these groups. We observed a significant increase in the number of distinct IGH clonotypes in samples with Hashimoto’s thyroiditis (p-value= 1.5×10^-5^) (**Figure S11**). The number of clonotypes varied from 113 for focal Hashimoto’s thyroiditis to 5621 for late-stage Hashimoto’s thyroiditis (**Figure 6a**). In addition, high clonotype diversity in kidney samples indicated the presence of glomerulosclerosis. In lung samples, high clonotype diversity corresponded to inflammatory diseases such as sarcoidosis and bronchopneumonia.

**Figure 6.**
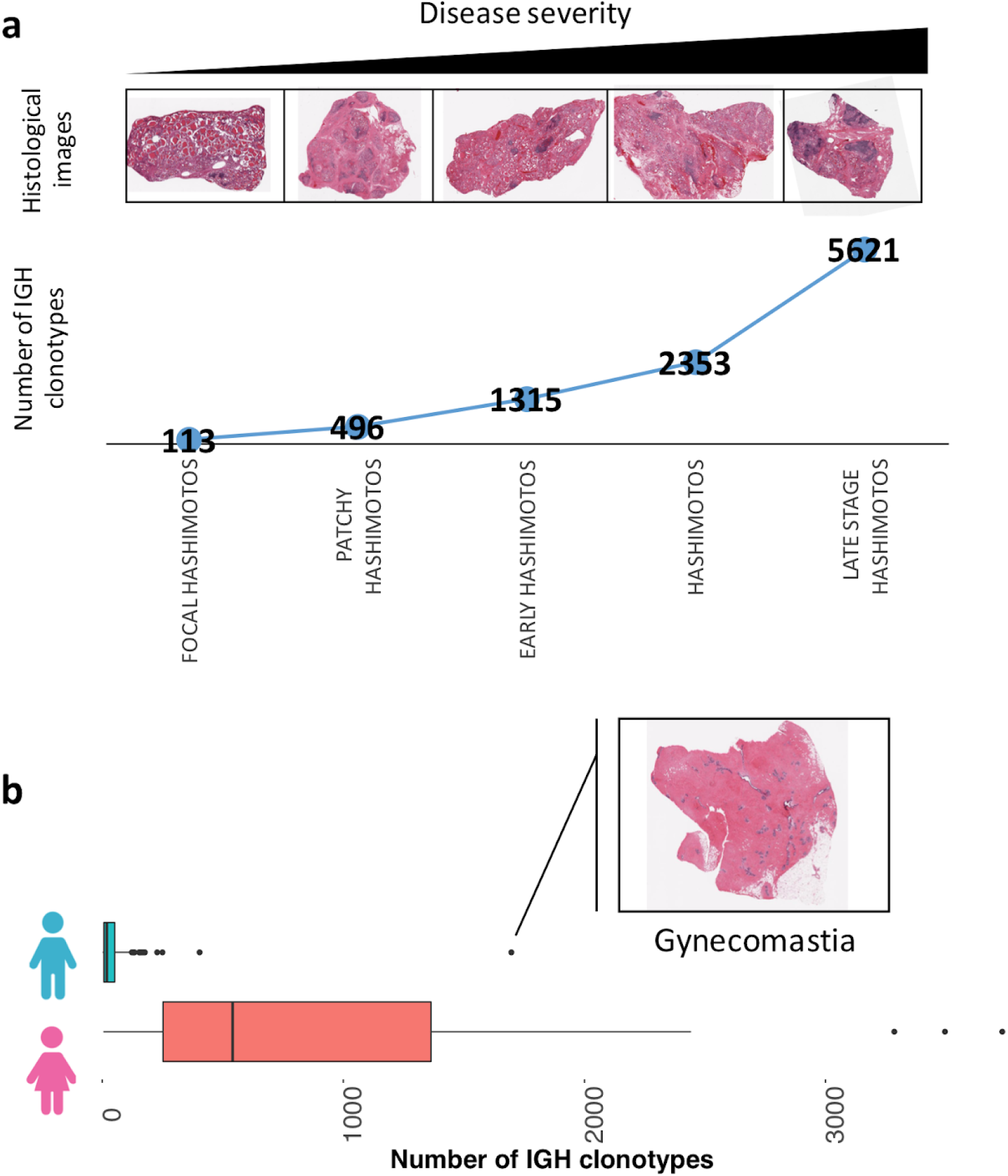
ImReP is able to identify samples with high activity of lymphocytes. Histological images of tissue cross-sections and pathologists’ notes have been used to validate the ImReP’s ability to detect the samples with a high activity of lymphocytes. **(a)** Samples were ordered by Hashimoto’s thyroiditis severity, as reported by pathologists’ notes. Histological images are provided to illustrate the disease state. The average number of Ig clonotypes is reported for each disease group. **(b**) Boxplot reporting number of clonotypes in the breast tissues for males and females. The outlier among the male samples is illustrated with the histological image.

We observed no difference in clonal diversity in males and females across the tissues, except in breast tissues (p-value<3.2×10^−12^). Increased clonotype diversity of breast tissue in male individuals corresponded to gynecomastia, a common disorder of non-cancerous enlargement of male breast tissue (**Figure 6b**).

## Discussion

We have developed a novel computational approach (ImReP) for reconstruction of Ig immune repertoires using RNA-Seq data. We demonstrate the ability of ImReP to efficiently extract Ig-derived reads from the RNA-Seq data and accurately assemble the corresponding hypervariable region sequence. The proposed algorithm can accurately assemble CDR3 sequences of Ig receptors despite the presence of sequencing errors and short read length. Simulations generated using various read lengths and coverage depth show that ImReP consistently outperforms existing methods in terms of precision and recall rates.

We have demonstrated the feasibility of applying RNA-Seq to study the adaptive immune repertoire. Although RNA-Seq lacks the sequencing depth of targeted sequencing (Rep-Seq), it can compensate by examining a larger sample size. Using ImReP, we have created the first systematic atlas of immunological sequence for Ig receptor repertories across diverse human tissues. This provides a rich resource for comparative analysis of a range of tissue types, most of which have not been studied before. The atlas of immune repertoires, available with the paper, is one the largest collection of CDR3 sequences and tissue types. We anticipate that this database will enhance future studies in areas such as immunology and contribute to the development of therapies for human diseases.

Using RNA-Seq to study immune repertoires has some advantages, including the ability to simultaneously capture clonotype populations from all the chains during a single run. It also allows simultaneous detection of overall transcriptional responses of the adaptive immune system, by comparing changes in the number of Ig transcripts to the much larger transcriptome. Given a large number of large-scale RNA-Seq datasets becoming available, we look forward to scaling up the atlas of immune receptors in order to provide valuable insights into immune responses across various autoimmune diseases, allergies, and cancers.

## Extended Experimental Procedures

For manuscript “**Profiling immunoglobulin repertoires across multiple human tissues by RNA Sequencing**”

### RNA-Sequencing data

We used RNA-Sequencing data from Genotype-Tissue Expression study (GTEx Consortium v.6) corresponding to 8,555 samples collected from 544 individuals from 53 tissues obtained from Genotype-Tissue Expression study (GTEx v6). RNA-Seq data is from Illumina HiSeq sequencing of 75 bp paired-end reads. The data was derived from 38 solid organ tissues, 11 brain subregions, whole blood, and three cell lines of postmortem donors. The collected samples are from adults matched for age across males and females.

### RNA-Sequencing data preprocessing

We downloaded the mapped and unmapped reads in BAM format from dbGap (http://www.ncbi.nlm.nih.gov/gap). For each, the sample we prepared the candidate receptor-derived reads to be the input for ImReP tool. First, we extracted reads mapped to the TCR and BCR genes. Some high throughput aligners allow partial mapping (soft clipping) resulting in cutting one or two ends of the reads and mapping the remaining read. Reads containing CDR3 sequences may be among those reads and are extracted by ImReP. The coordinates of TCR and BCR genes (GRCh37 human reference genome release) are provided Table S3. Second, we prepared the high-quality unmapped reads using ROP (step1 – step3) https://sergheimangul.wordpress.com/rop/ (Mangul et al.) by filtering out low quality, low complexity reads and reads that match rRNA repeats. We also filtered out lost human reads (reads unmapped to the reference genome) and lost repeat reads (unmapped reads mapped to the repeat sequences). The reads mapped to the BCR and TCR loci and high-quality unmapped reads were merged, and ImReP used this data to assemble CDR3 sequences and corresponding V(D)J recombinations.

### ImReP algorithm

ImReP is a computational approach to assemble CDR3 sequences and detect corresponding V(D)J recombinations from B and T cell receptors. ImReP is composed of two stages. In the first stage, ImReP utilizes the reads that simultaneously overlap V and J gene segments to infer the CDR3 sequences. We define the CDR3 as the sequence of amino acids between the cysteine on the right and phenylalanine (for TCR, IGK, and IGL) or tryptophan (for IGH) on the left of the junction. We first convert the read sequences from nucleotides to amino acids. We scan the *amino acid* sequences of the read and determine the putative CDR3 as a subsequence of the read starting from cysteine (C) and ending with phenylalanine (F) (and tryptophan (W) for IGH). The reads containing the described substring are considered candidate CDR3 reads. We denoted n to be the length of the read. We denoted the coordinates of putative CDR3 string to be x and y, corresponding with the start and end of the CDR3 sequence in the read coordinates. This way each candidate CDR3 read is composed of three parts:

1. r[0,x-1] is a prefix of the read, potentially overlapping suffix of V gene. It contains the amino acids from the read, from position 0 to x-1.
2. r[x,y] is a substring of the read containing the putative CDR3 sequence. It contains the amino acids from the read, from position x to y.
3. r[y+1,n] is a suffix of the read potentially overlapping prefix of J gene. It contains the amino acids of the read, from position y+1 to n

The amino acid sequences of V and J genes of B cell receptors (BCR) and T cell receptors (TCR) were imported from IMGT (International ImMunoGeneTics information system) (http://www.imgt.org/vquest/refseqh.html#V-D-J-C-sets). For each V gene, we identify last conserved cysteine (C) and record its position pC. For each J gene, we identify first conserved phenylalanine (for IGK, IGL, TCRA, TCRB, TCRG, TCRD) or tryptophan (for IGH) and record its position pF. For each V gene, we extract two substrings: Vx=V[0, p_C_-1] and Vy=[p_C_+1,nV]. For each J gene, we record two substrings: Jx=J[0, p_F_-1] and Jy=J[p_F_+1,nJ], where nV and nJ are the lengths of V and J genes, respectively. Given a set of candidate CDR3 reads, we attempt to find the corresponding V and J genes. We match a substring of the read r[0,x-1] with the corresponding suffix of Vx for V genes. We also match the read r[y+1,n] with the corresponding prefix of Jx for J genes. We consider a read to match V gene if the length of r[0,x-1] is greater than 4 and the edit distance between r[0,x-1] and Vx is less than 2. We consider a read to match J gene if the length of r[y+1,n] is greater than 4 and the edit distance between r[y+1,n] and Jx is less than 2. In case a read overlaps equally (in terms of edit distance) among multiple V genes and J genes, we report all of them.

In the second stage, ImReP utilized the reads overlapping only V or J gene. Such reads contain a partial CDR3 sequence. ImReP builds a suffix tree S on the reads overlapping any of the V genes. Then, for each read j overlapping a J gene a V-gene overlapping read, v from S is determined (in case if any exists). Reads v and j are concatenated (with overlap), and the CDR3 region is extracted.

Further, ImReP uses a CAST clustering technique to correctly assemble CDR3s for PCR and sequencing errors. The output of the algorithm is the set of CDR3 partitions, and each of the partition corresponds to a clonotype. Specifically, ImReP builds a complete graph G = (V, E, w), where the set of vertices V is represented by the set of assembled CDR3 sequences. The weight of the edge is determined by the inverse of the edit distance, computed between the two CDR3 sequences x and y. The CAST algorithm is executed with the following procedure. A new partition P is initialized with the max-degree node. Then, the set of jjclose’’ vertices is iteratively added to the partition, and the set of jjdistant’’ vertices are removed from the partition. A vertex v is deemed to be jjclosejj (jjdistant’’), if the average distance from v to the vertices from P is greater (smaller) than a user-defined threshold. The procedure is repeated until either the set of jjclose’’ or the set of jjdistant’’ vertices is empty. In such a way, the partition P is based on a max-degree node and extended with the jjclose’’ vertices. Vertices belonging to P are then removed from the graph G, and the clustering procedure is repeated until all of the vertices are assigned to a partition. Let {v_1_,v_2_,…,v_i_,…,v_n_} be a partition output by the CAST algorithm. Each v_i_ has an associated weight equal to the count of CDR3’s v_i_, which was assembled during the first two stages of ImReP. We compute the weighted consensus sequence of P and output the sequence as a final clonotype. Finally, we map D genes (for IGH, TCRB, TCRG) onto assembled CDR3 sequences and infer corresponding V(D)J recombination. Starting with release v0.8, ImReP reports out of frame CDR3 sequences.

### Validation based on simulated RNA-Seq data

We performed in-silico simulations to investigate the feasibility of using RNA-Seq to study the clonal adaptive immune repertoire. We first checked the ability of the ImReP to extract the receptor-derived reads from the RNA-Seq reads. First, we simulated the TCR and BCR transcripts, which are composed of recombined VDJ segment with non-template insertion at the V(D)J junction (Figure S2). We used the IMGT database (http://www.imgt.org/vquest/refseqh.html) of V and J gene segments. We randomly selected V, D and J segments and inserted a sequence of random nucleotides between V and D and between D and J. The length of the inserted sequence was sampled from the Gaussian-like distribution with mean 15 (Miqueu, Patrick, et al.). We also exclude the simulated transcripts with the random insertions leading to out-of-frame proteins. We used LymAnalizer (https://sourceforge.net/projects/lymanalyzer/) validate CDR3 sequences of the transcript. We used SimNGS (https://www.ebi.ac.uk/goldman-srv/simNGS/) to simulate paired-end reads from BCR and TCR transcripts referred as receptor-derived reads. Next, we simulated 50 million transcriptomic reads from the human transcriptome reference (GRCh37). We mixed receptor-derived reads with transcriptomic reads into an RNA-Seq mixture (Figure S3). We then applied ImReP to a simulated RNA-Seq mixture to check the ability of ImReP to extract CDR3-derived reads from the RNA-Seq mixture.

Next, we studied the effects of the coverage and read length on the ability to reconstruct CDR3 sequences. In total, we simulated 1,000 BCR or TCR transcripts. We simulated paired-end reads of various read length (l=50,75,100) with use various coverage of TCR and BCR transcripts (c=1, 2, 4, 8, 16, 32, 64, 128). We used the power law distribution to assign frequencies to simulated T and B cell transcripts (Weinstein, Joshua A., et al.). The CDR3 amino acid sequences assembled by ImReP were compared to simulated transcripts to evaluate the recall and precision for various read length and coverage (main text and Figure 2).

We define recall and precision in the following way:

- Recall = TP/(TP+FN)
- Precision = TP/(TP+FP)

Where TP is the number of correctly assembled CDR3 sequence features (exact match to the simulated CDR3), FN is the number of simulated CDR3 sequence features not assembled by the method, and FP is the number of incorrectly assembled CDR3 sequences. Scripts to simulate BCR and TCR transcripts and the reads are available online at https://smangul1.github.io/ImReP-Gtex/

### Comparison to other methods

We compared the ImReP to existing methods based on simulated data generated, as described in “Validation based on simulated RNA-Seq data”. We compared ImReP to the following tools:

⋅ TRUST, https://bitbucket.org/lukeli1987/trustnew
⋅ MIXCR, http://mixcr.readthedocs.io/en/latest/
⋅ IMSEQ, http://www.imtools.org/
⋅ IGBLAST pipeline, https://github.com/nbstrauli/influenza_vaccination_project
⋅ V’DJer, https://github.com/mozack/vdjer
⋅ TraCeR, https://github.com/Teichlab/tracer

IMSEQ cannot be applied to RNA-Seq reads, because they were originally designed for Rep-Seq, targeted sequencing of BCR or TCR loci. Thus, it was provided with only the simulated receptor-derived reads to assess accuracy.

We used the following command to run MiXCR in the RNA-Seq mode with the option to filter *out-of-frame* CDR3 sequences (version 2.2 of MiXCR was used):

⋅ mixcr align -p rna-seq -s hsa -r log.txt r1.fastq r2.fastq alignments.vdjca -c $chain
⋅ mixcr assemblePartial -r log.txt -p alignments.vdjca alignments_rescued_1.vdjca
⋅ mixcr assemblePartial -r log.txt alignments_rescued_1.vdjca alignments_rescued_2.vdjca
⋅ mixcr assemble -r log.txt alignments_rescued_2.vdjca clones.clns
⋅ mixcr exportClones clones.clns clones.txt

The following command was used to run IgBlast-based pipeline:

⋅ Bash pipeline.bash [path_to_input_directory] [path_to_output_directory] [path_to_IgBLAST_directory]

We run TRUST in single-end and paired-end mode.

- trust-default (considers paired-end information) TRUST.py -f <bam> -a
- trust-se (treats reads as single-end reads) TRUST.py -f <bam> -a -s

We run IMSEQ with default parameters for IGH, IGK, IGK,TCRA, and TCRB chains using the following commands:

⋅ imseq -ref Homo.Sapiens.TRA.fa -o output-file_TCRA.tsv <fastq>
⋅ imseq -ref Homo.Sapiens.TRB.fa -o output-file_TCRB.tsv <fastq>
⋅ imseq -ref Homo.Sapiens.IGH.fa -o output-file_IGH.tsv <fastq>
⋅ imseq -ref Homo.Sapiens.IGK.fa -o output-file_IGK.tsv <fastq>
⋅ imseq -ref Homo.Sapiens.IGL.fa -o output-file_IGL.tsv <fastq>

Where <fastq> is a file with the unmapped reads plus reads mapped to BR and TCR loci.

We run VDJer using the following command:

⋅ vdjer --in $bam --ins 175 --chain IGH --ref-dir igh
⋅ vdjer --in $bam --ins 175 --chain IGK --ref-dir igk
⋅ vdjer --in $bam --ins 175 --chain IGL --ref-dir igk

Where ’$bam’ contains mapped and unmapped reads

We were not able to run VDJer on both real and simulated reads. We reported the issues here:

⋅ https://github.com/mozack/vdjer/issues/7
⋅ https://github.com/mozack/vdjer/issues/8

The following command were used to run TraCeR (v.0.4.0)

⋅ tracer assemble -c tracer.conf -p 4 -s Hsap r1.fastq r2.fastq cellname tracer_output We have prepared the configuration file for TraCeR according to the software recommendations. Tracer doesn’t work with Bowtie2 version 2.3.0. It has to be 2.2.9 or older.

Scripts and commands utilized to repertoire assembly tools are available online at https://smangul1.github.io/ImReP-Gtex/

### Cell type composition

B cell signature values per sample were derived from the SaVant signature visualization tool (Lopez et al.). Cell-specific signature genes are first defined from a set of cells/tissues obtained from the Human Body Atlas (Su AI et al., 2004) by using the proportional median values. We calculate these values by dividing the intensity of a probe in a particular cell type by its median value across all cells/tissues. The top 25 genes with the highest proportional median value for CD19+ B cells were defined as the specific signature for that cell type (**Table S5**). Any BCR genes were removed from the signature. The signature score is then generated from the average of the log2-transformed values of the signature genes within each sample.

### Definition of clonotype

Clonotypes are defined as clones with identical CDR3 amino acid sequences.

### Histological images and pathologist notes

We used histological images and pathologists’ notes (available at GTEx portal, http://www.gtexportal.org/home/histologyPage#data) to validate the adaptive immune profile of the samples. Although samples were derived from primary tissues, they often have a mixed cell type composition. For example, samples from stomach tissues have various proportions of lymphocytes as they were derived from mucosal or muscularis part of the tissue (based on pathologists’ notes). GTEx samples with inflammation and/or subject to various diseases are investigated separately. Pathologists’ notes report the percentage of mucosa and disease or inflammation status of the tissue.

### Software Availability

ImReP (https://github.com/mandricigor/imrep/wiki) is freely available as source code. It takes mapped RNA-Seq reads as bam or sam, and it assembles *Ig* clonotypes and corresponding V(D)J recombinations. ImReP is distributed under the terms of the General Public License version 3.0 (GPLv3).

### Data representation

We have used WebLogo3 (http://weblogo.threeplusone.com/manual.html) to represent the amino acid composition of assembled CDR3 sequences.

### Data Availability Statement

The RNA-Seq data discussed in this paper is available as part of the Genotype-Tissue Expression (*GTEx*) Project.

---

**Figure S1.**
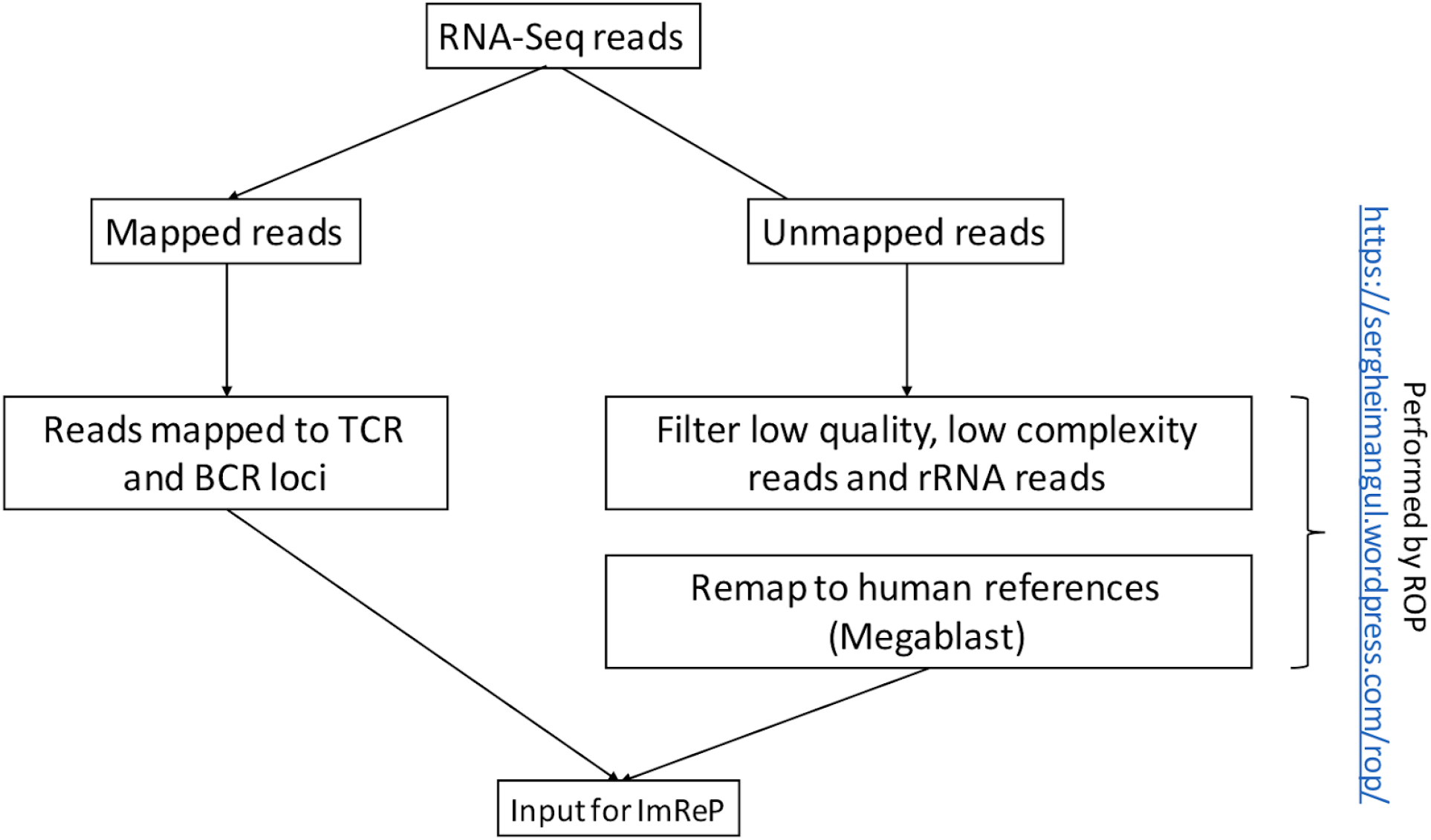
Schematic on how to select the candidate receptor-derived reads from RNA-Seq reads, which are the input for ImReP.

**Fig S2.**
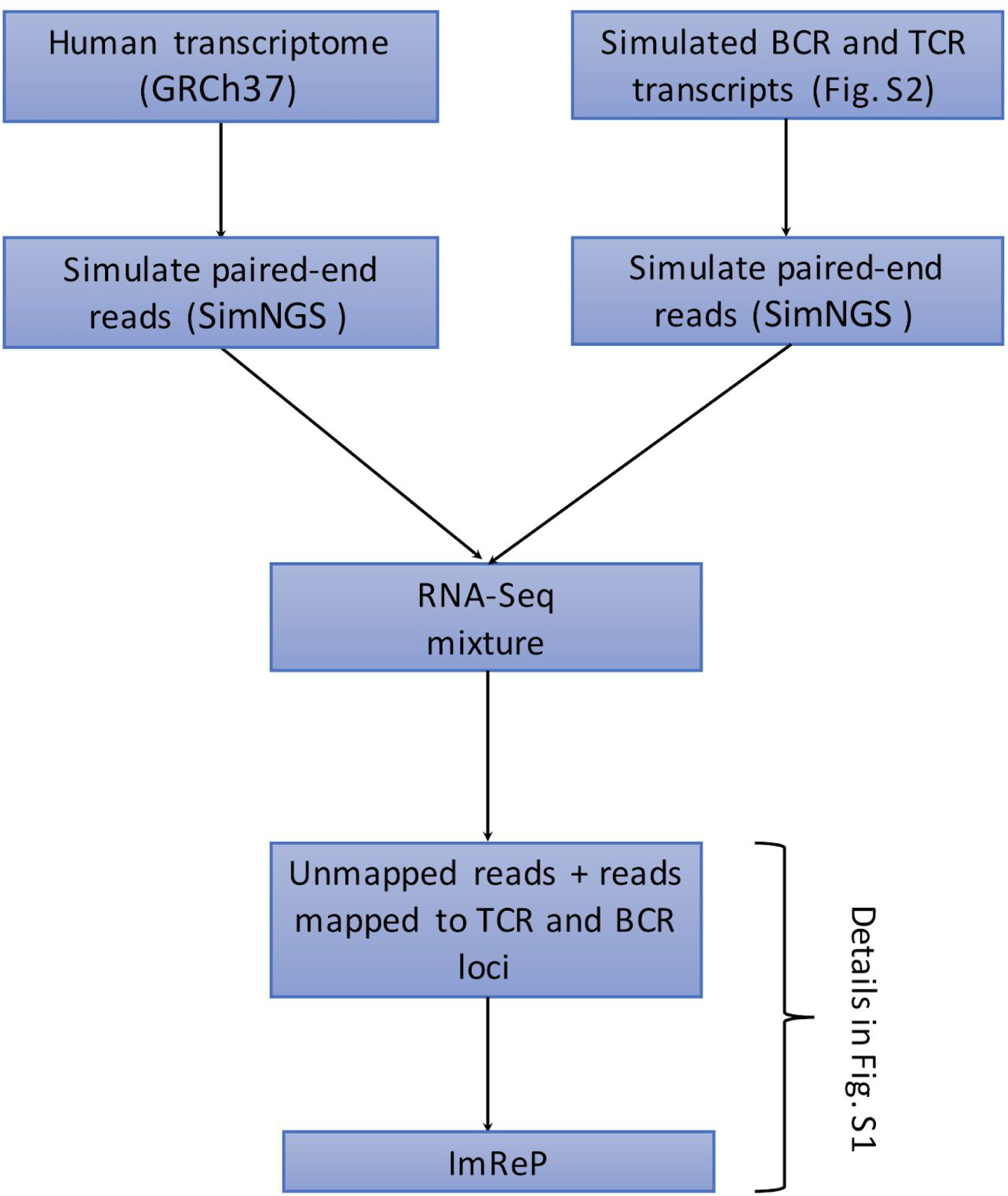
Workflow of BCR and TCR transcript simulations.

**Figure S3.**
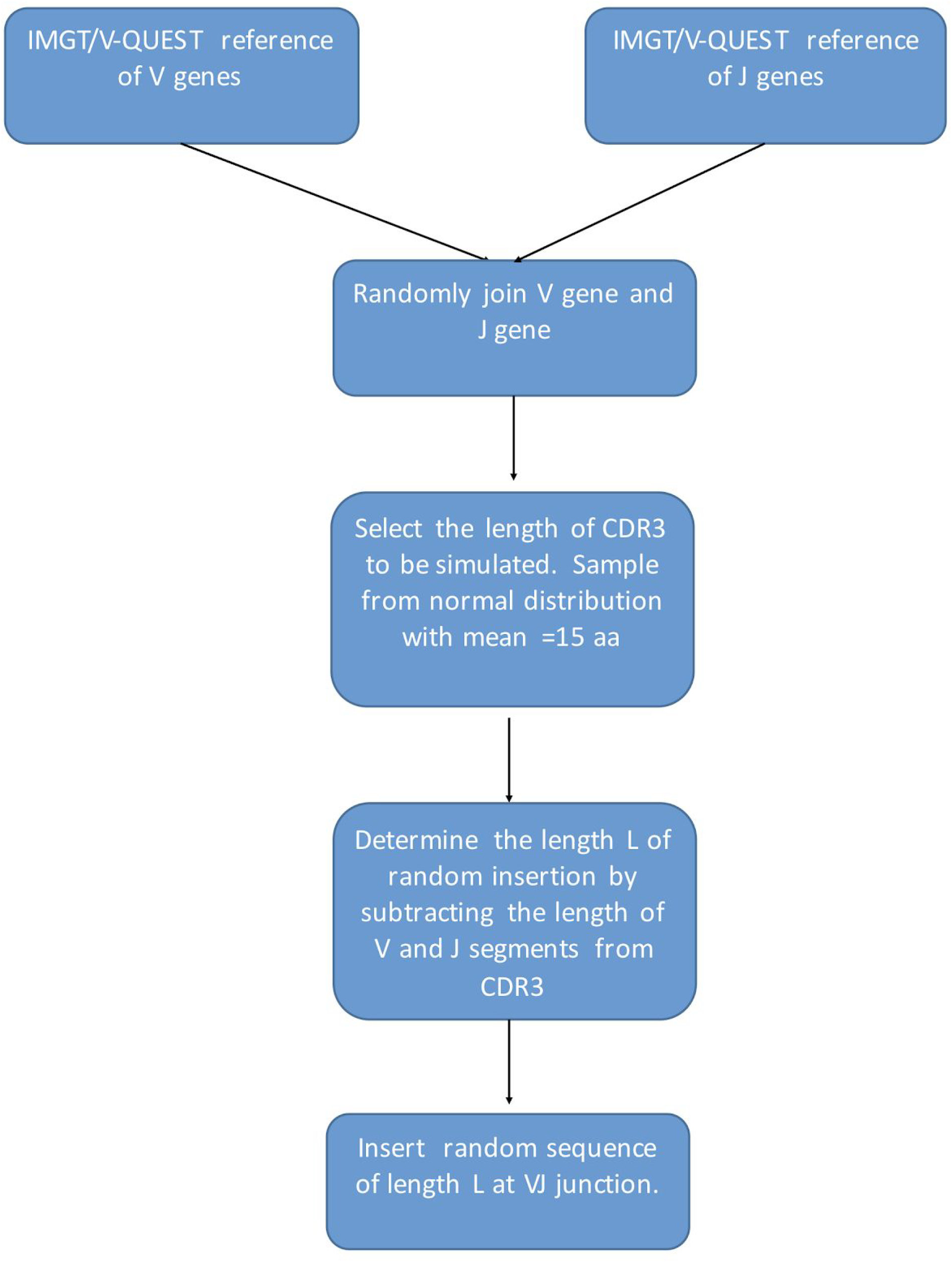
Schematic on how the mixture of transcriptomic and receptor-derived reads was generated.

**Figure S4.**
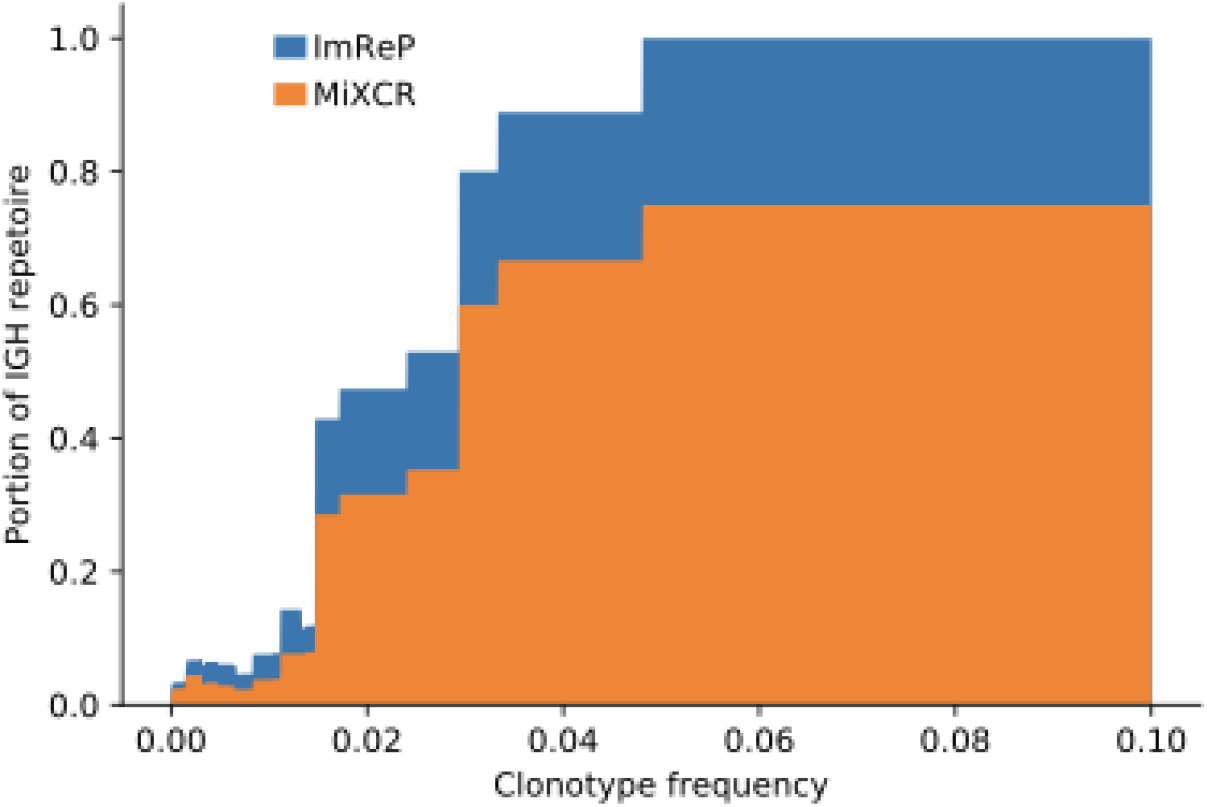
Concordance of targeted BCR-Seq and non-specific RNA-Seq performed on 13 tumor biopsies from Burkitt lymphoma. Area chart shows the proportion of the total IGH repertoire captured by ImRep (blue) and MiXCR (RNA-Seq mode) (orange) depending on the minimum BCR-seq-confirmed clonotypes frequency considered. The *x-axis* corresponds to BCR-seq-confirmed clonotypes frequency Z. The *y*-axis corresponds to the fraction of assembled IGH repertoire with clonotype abundances greater than Z. The total repertoire was defined as the total number of the BCR-seq-confirmed clonotypes.

**Figure S5.**
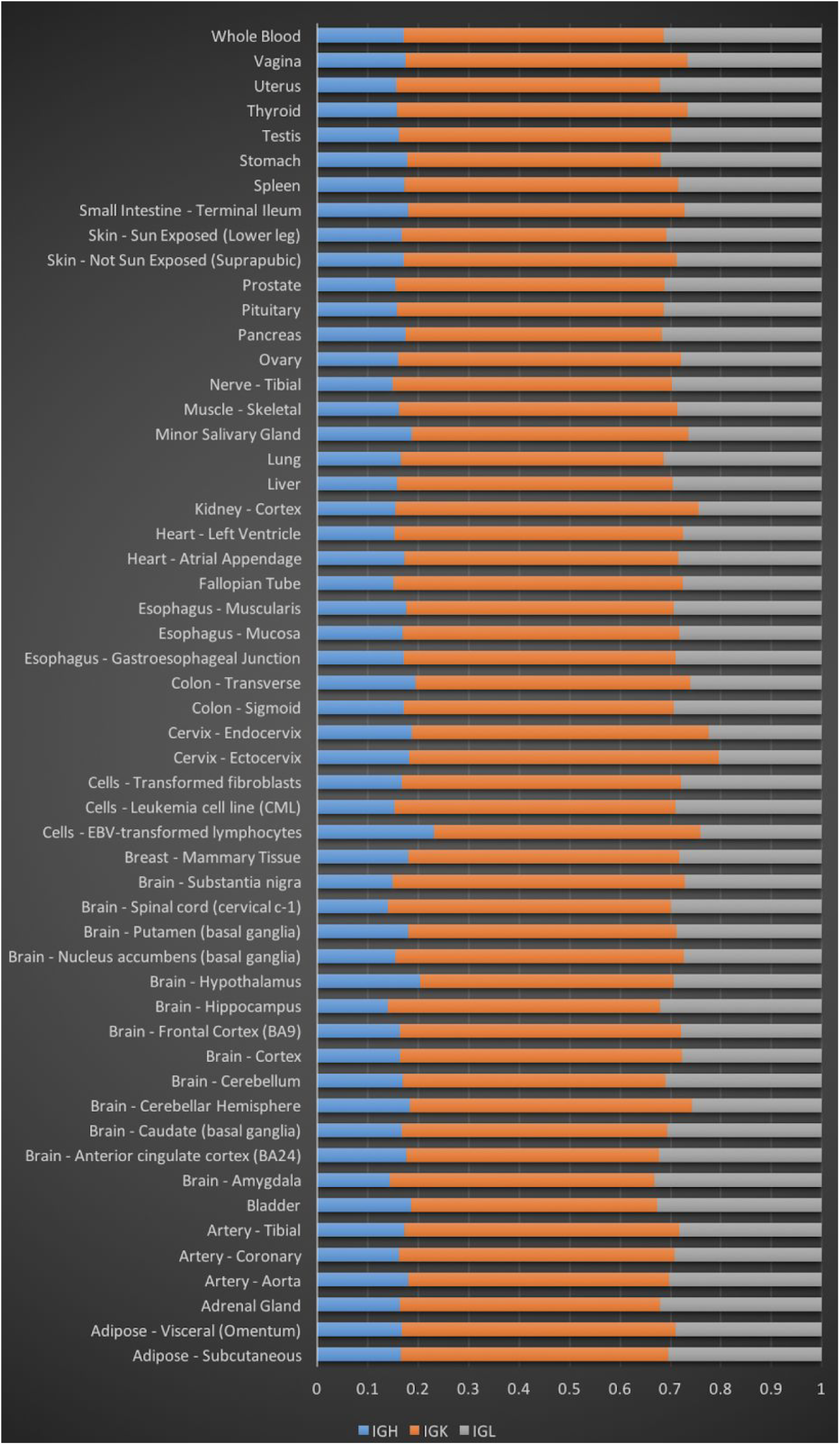
The fraction of IGH, IGK, and IGL among the whole B cell population across 53 body sites. The fraction is calculated based on the number of reads derived from CDR3 sequences of each BCR chain.

**Figure S6.**
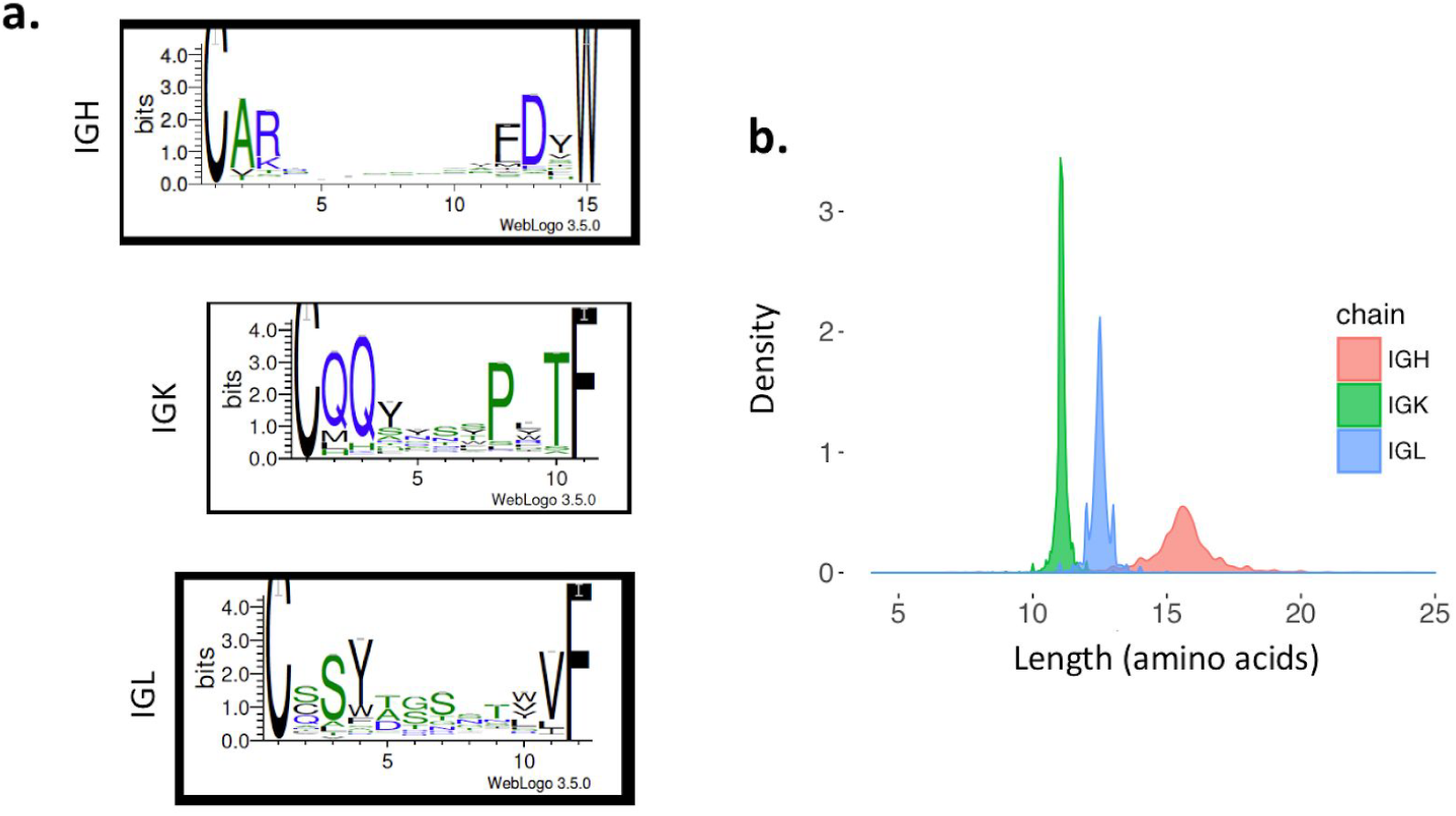
The length and amino acid composition of the assembled CDR3 sequences of immunoglobulin receptor chains. The sequence logo (using WebLogo) of amino acid composition representation for CDR3 sequences of mean length. The height of the amino acid within the stack indicates the relative frequency. Distribution of CDR3 sequence length is estimated using kernel density. (a) Sequence logo of a 15-amino-acid CDR3 sequence of IGH. (b) Sequence logo of 11-amino-acid CDR3 of IGK. (c) Sequence logo of a 12-amino-acid CDR3 sequence of IGL. (h) Distribution of CDR3 sequence length is estimated using s kernel density separately for each Ig chain.

**Figure S7a.**
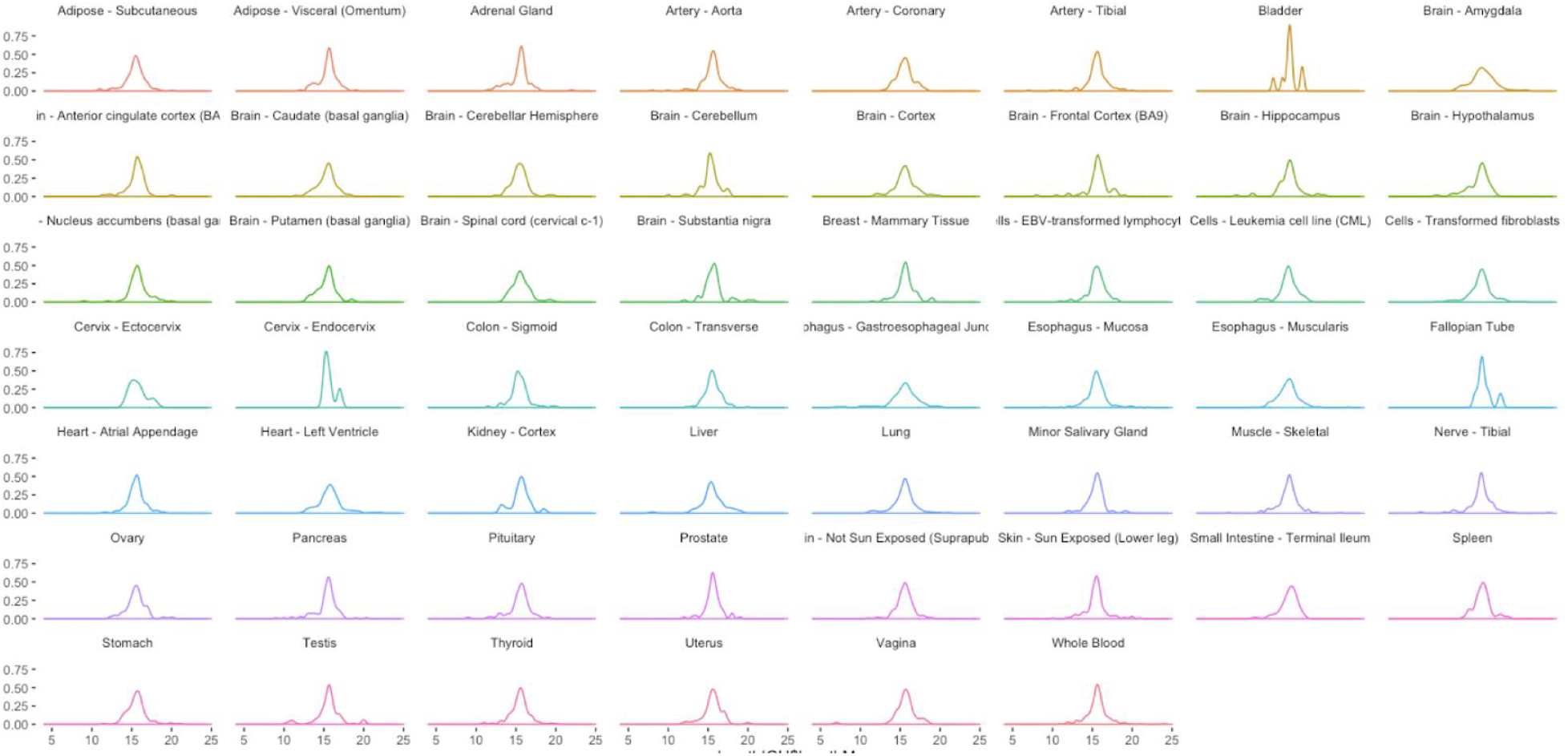
Length distribution of amino acid sequences of the CDR3 region in immunoglobulin heavy chain (IGH), presented across 53 various body sites.

**Figure S7b.**
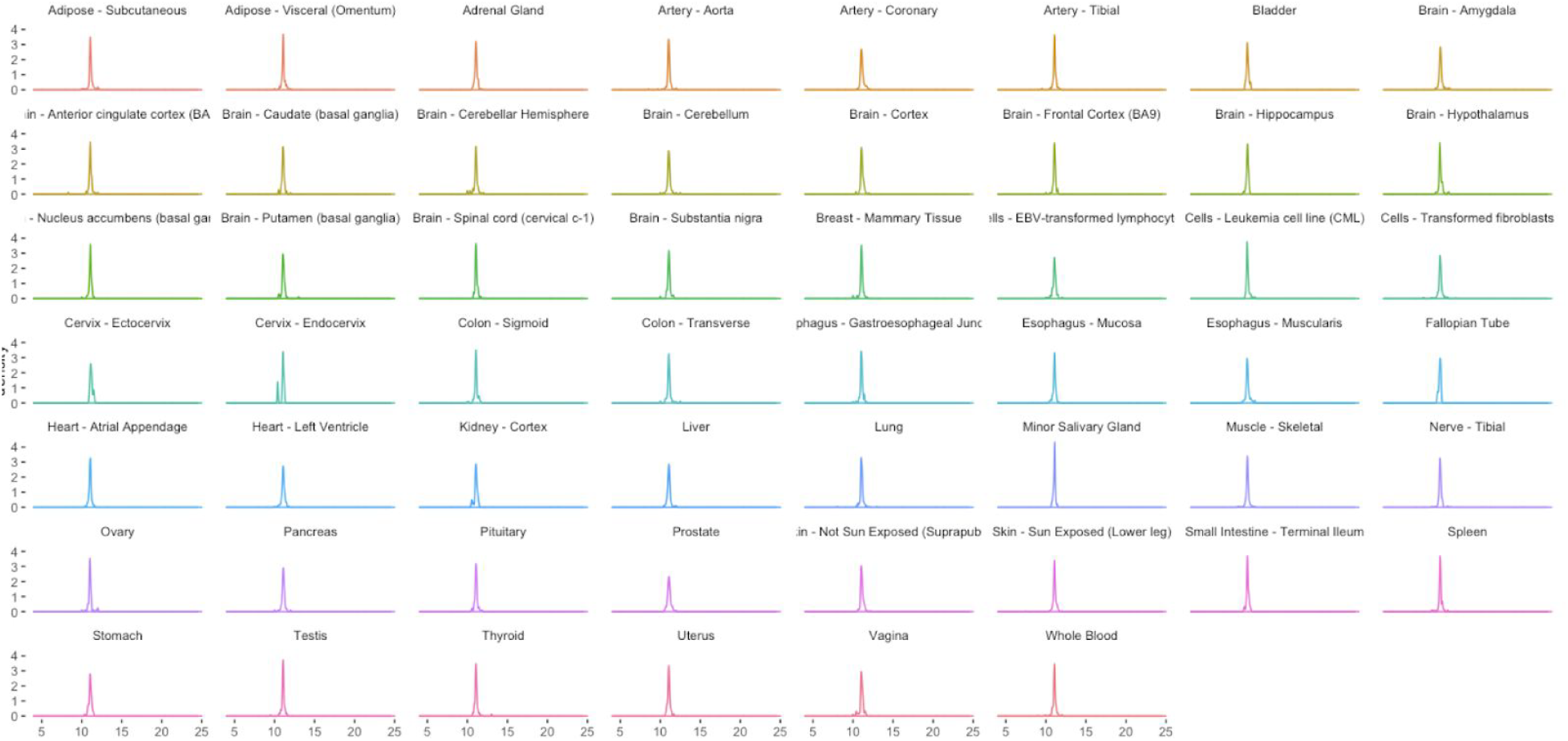
Length distribution of amino acid sequences of the CDR3 region in immunoglobulin kappa chain (IGK), presented across 53 various body sites.

**Figure S7c.**
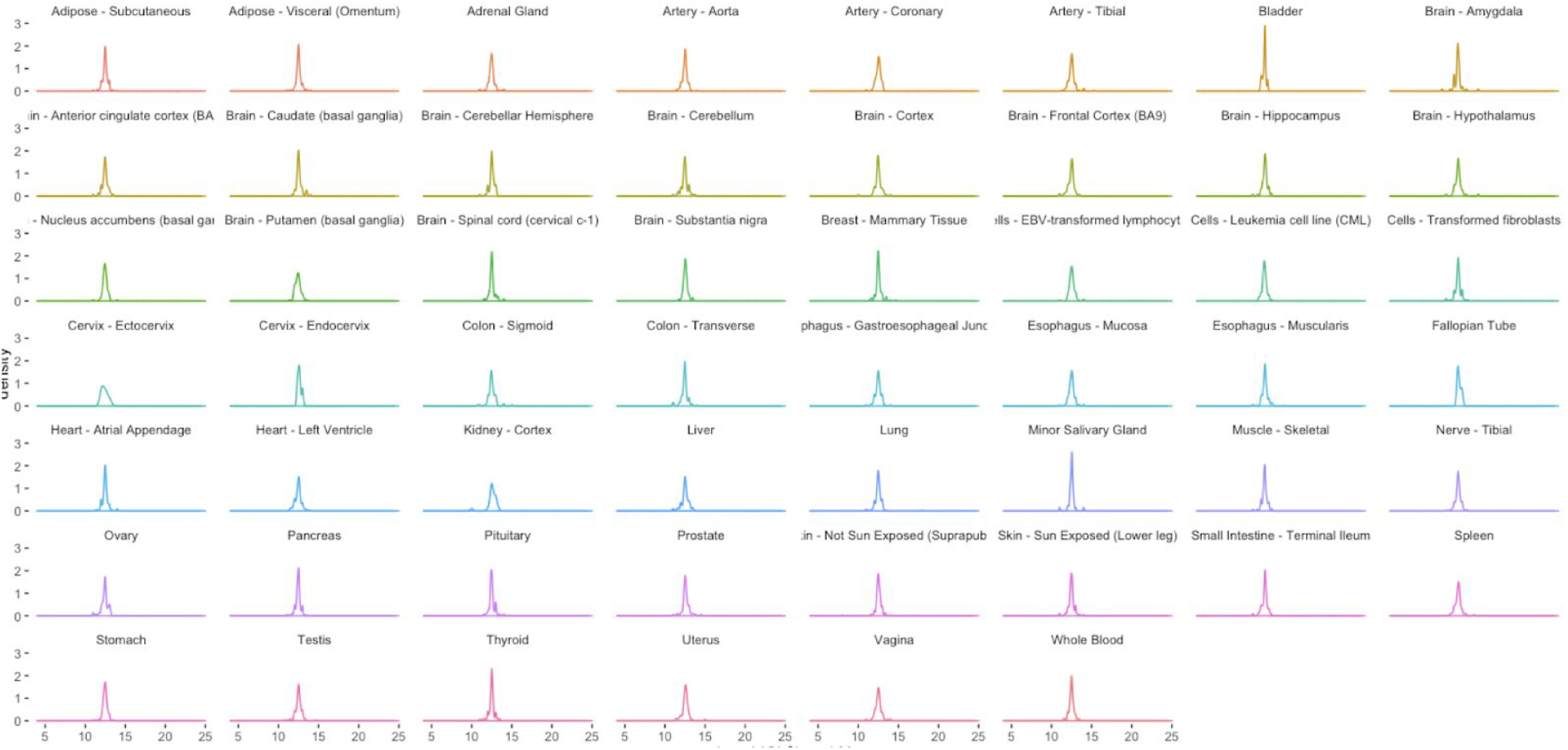
Length distribution of amino acid sequences of the CDR3 region in immunoglobulin kappa chain (IGK), presented across 53 various body sites.

**Figure S8.**
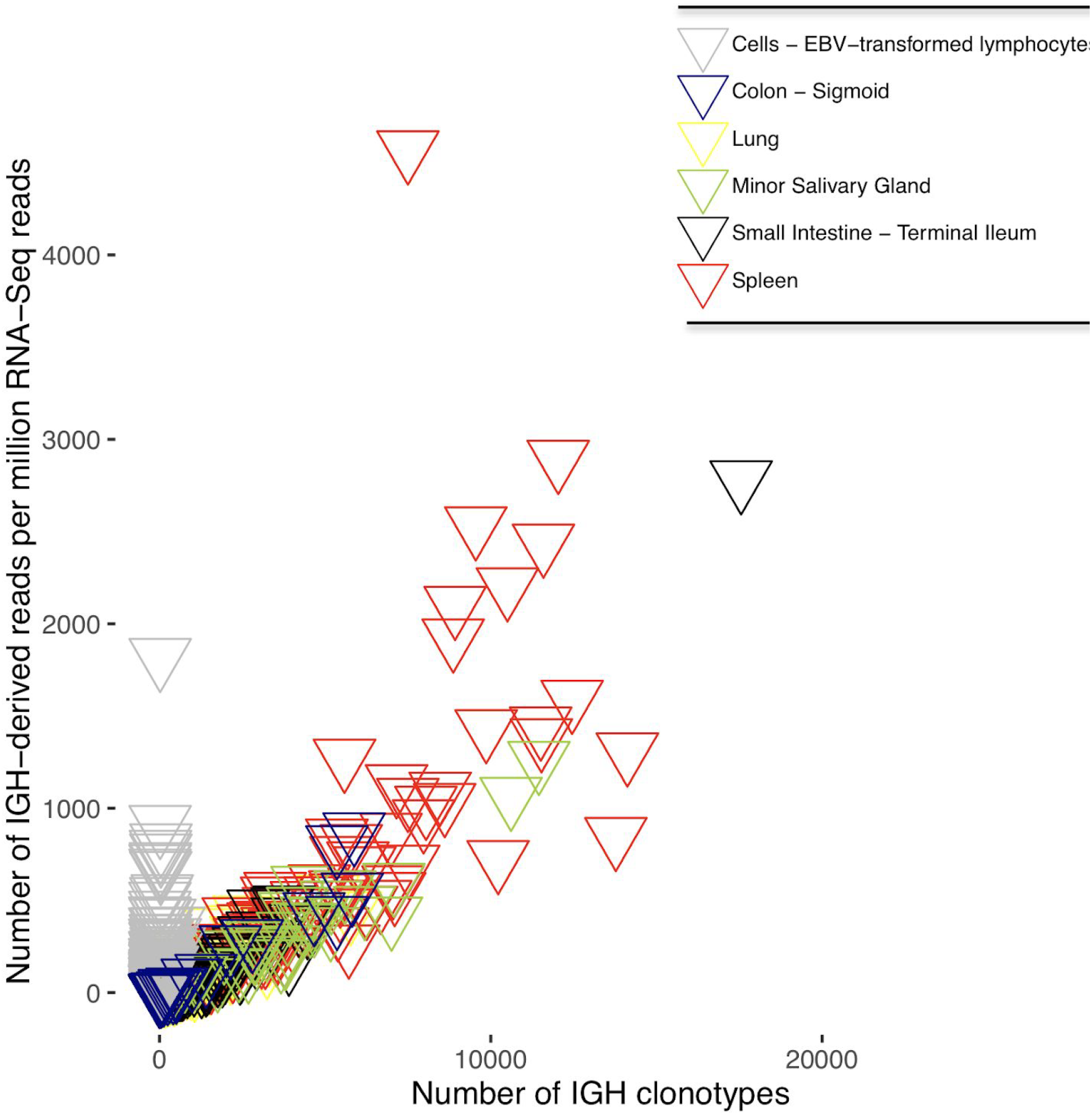
Scatterplot of the number of IGH clonotypes (CDR3s) in each sample, plotted against the number of IGH-derived reads per 1 million RNA-Seq reads.

**Figure S9.**
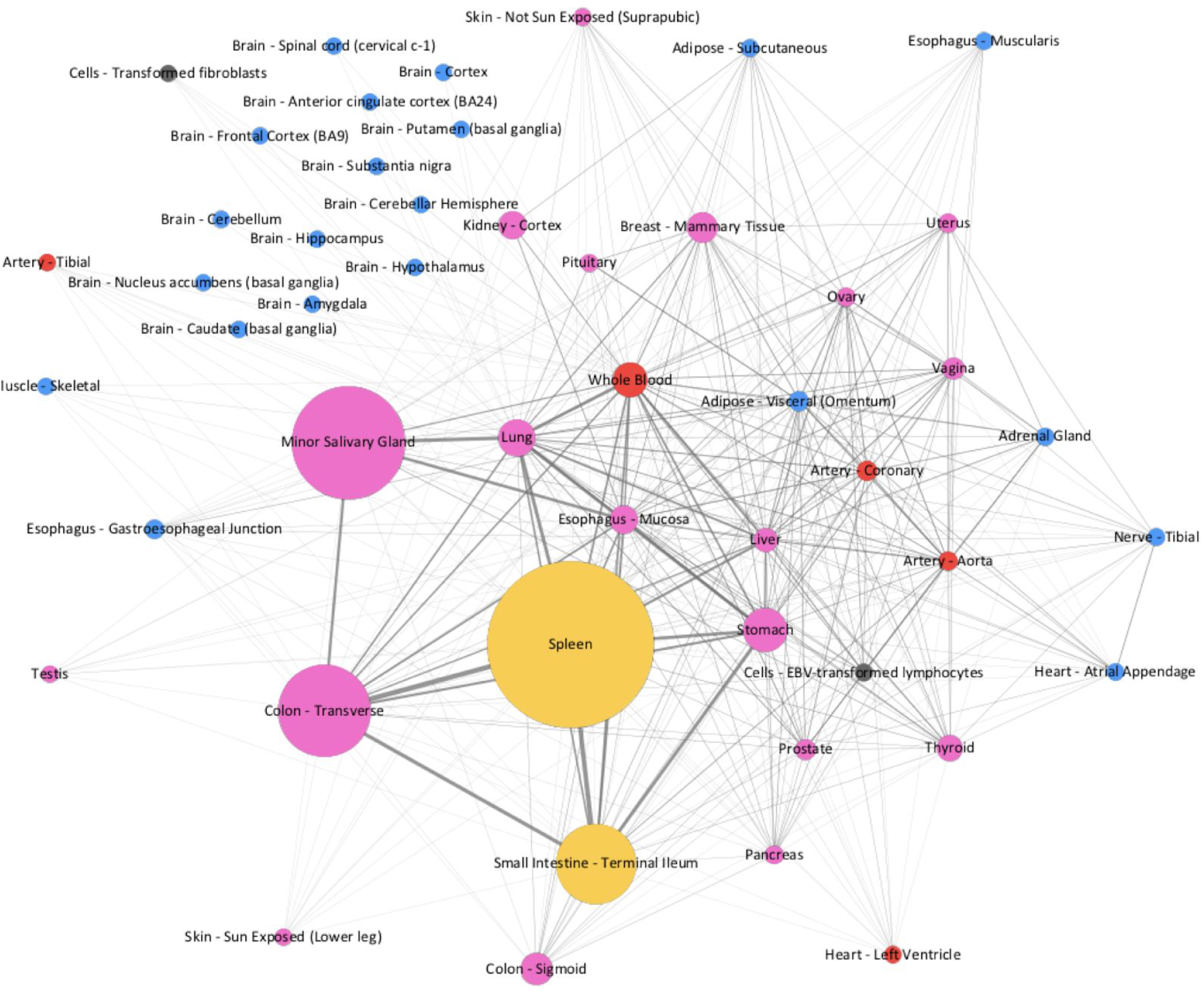
The flow of IGK clonotypes across diverse human tissues is presented as a network. Edges with beta diversity >.001 are presented.

**Figure S10.**
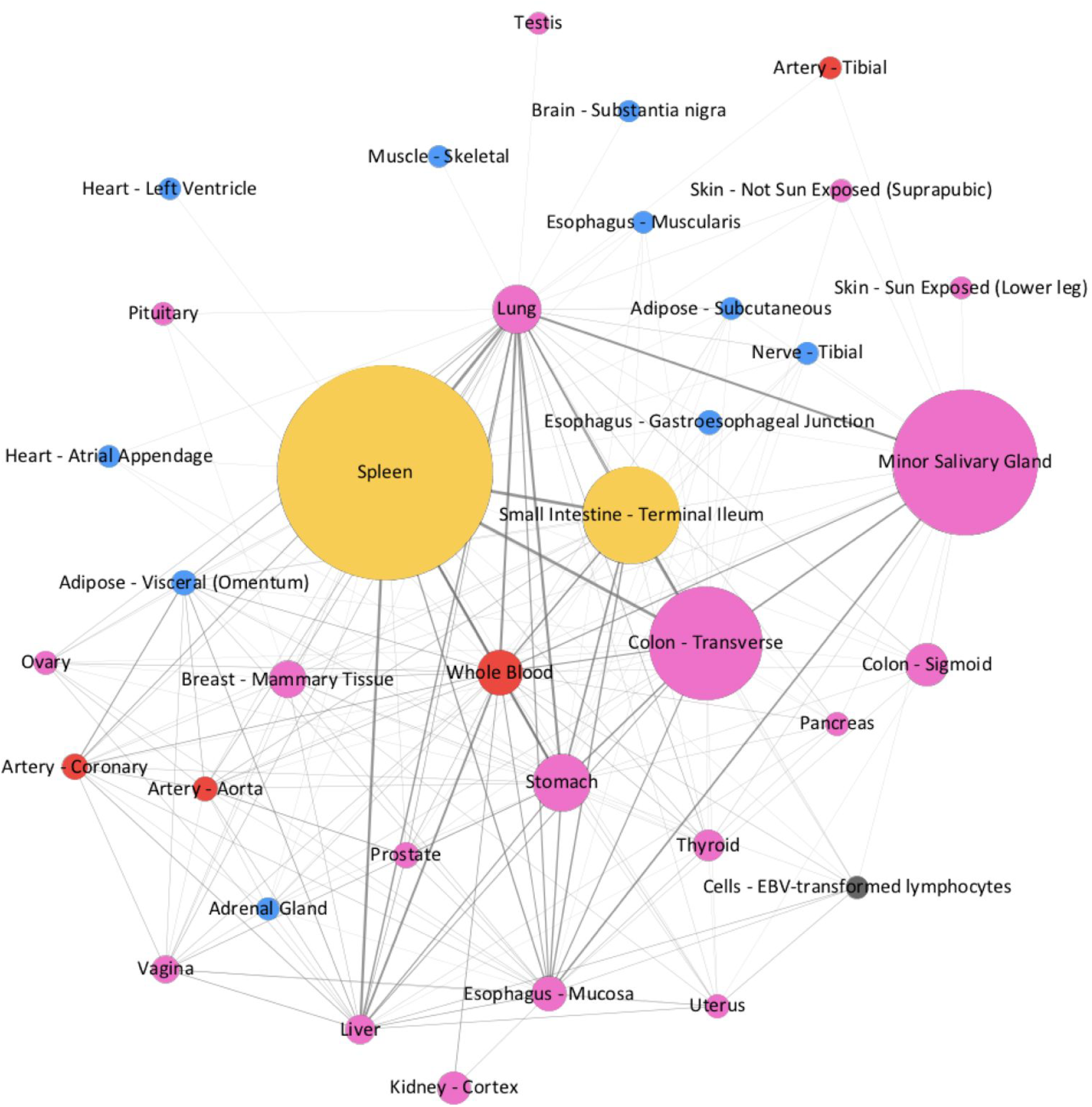
The flow of IGL clonotypes across diverse human tissues is presented as a network. Edges with beta diversity >.001 are presented.

**Figure S11.**
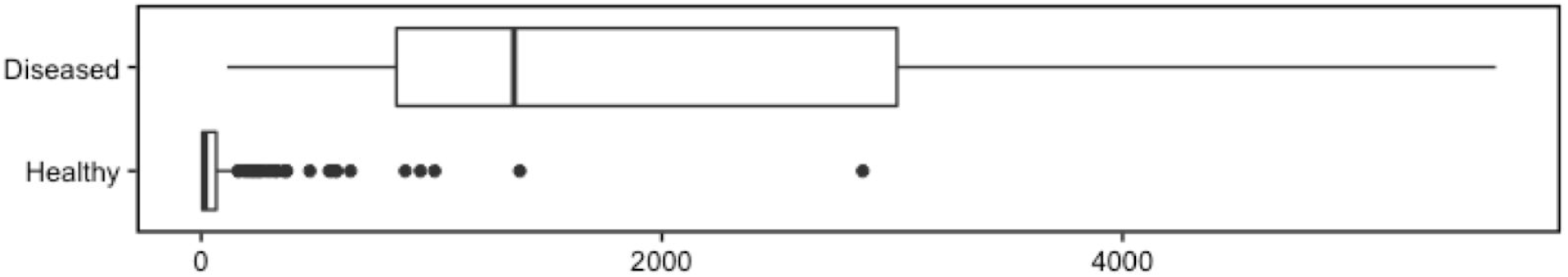
The number of IGH clonotypes for healthy individuals (Healthy) and individuals bearing Hashimoto’s disease (Diseased). Pathologists’ notes were used to annotate samples as healthy or diseased. A significant increase in the number of distinct IGH clonotypes in samples with Hashimoto’s thyroiditis (p-value= 1.5×10^−5^) is observed.

**Table S1.** Concordance of targeted BCR-Seq and non-specific RNA-Seq performed on 13 tumor biopsies from Burkitt lymphoma.

| ID | Major clonotype frequency (BCR-Seq) | Minor clonotype frequency (BCR-Seq) | Portion of IGH repertoire captured by ImRep | Portion of IGH repertoire captured by MiXCR | Major clonotype frequency (ImRep) | Minor clonotype frequency (ImRep) | Major clonotype frequency (MiXCR) | Minor clonotype frequency (MiXCR) | Number of BCR-SEQ-confirmed clonotypes | Number of ImRep-derived clonotypes (RNA-Seq) | Number of MiXCR-derived clonotypes (RNA-Seq) |
| --- | --- | --- | --- | --- | --- | --- | --- | --- | --- | --- | --- |
| 009-0192 | 0.64% | 0.16% | 7.48% | 7.96% | 0.64% | 0.16% | 0.64% | 0.16% | 583 | 34 | 32 |
| 009-0184 | 1.11% | 1.11% | 1.11% | 0.00% | 1.11% | 1.11% | 0.00% | 0.00% | 90 | 1 | 0 |
| 009-0148 | 1.45% | 0.15% | 0.00% | 0.00% | 0.00% | 0.00% | 0.00% | 0.00% | 671 | 0 | 0 |
| 009-0171 | 2.94% | 1.47% | 2.94% | 0.00% | 2.94% | 2.94% | 0.00% | 0.00% | 61 | 1 | 0 |
| 009-0203 | 3.33% | 0.83% | 0.00% | 0.00% | 0.00% | 0.00% | 0.00% | 0.00% | 117 | 0 | 0 |
| 009-0186 | 16.67% | 0.67% | 28.67% | 20.00% | 16.67% | 0.67% | 16.67% | 0.67% | 113 | 15 | 5 |
| 009-0174 | 62.54% | 0.34% | 62.54% | 0.00% | 62.54% | 62.54% | 0.00% | 0.00% | 55 | 1 | 0 |
| 009-0109 | 96.66% | 0.01% | 96.70% | 0.01% | 96.66% | 0.01% | 0.01% | 0.01% | 223 | 4 | 1 |
| 009-0249 | 97.68% | 0.00% | 97.68% | 97.68% | 97.68% | 97.68% | 97.68% | 97.68% | 379 | 1 | 1 |
| 009-0112 | 97.75% | 0.00% | 97.75% | 97.75% | 97.75% | 97.75% | 97.75% | 97.75% | 201 | 1 | 1 |
| 009-0122 | 98.16% | 0.00% | 98.17% | 98.16% | 98.16% | 0.00% | 98.16% | 0.00% | 119 | 3 | 2 |
| 009-02 | 99.72% | 0.00% | 99.72% | 99.75% | 99.72% | 99.72% | 99.72% | 99.72% | 50 | 1 | 2 |
| 02 |  |  |  |  |  |  |  |  |  |  |  |
| 009-01<br>03 | 99.75% | 0.03% | 99.75% | 99.75% | 99.75% | 99.75% | 99.75% | 99.75% | 9 | 1 | 1 |

**Table S2.** Data overview. Characteristics of 8,555 samples across 544 individuals from 53 body sites obtained from Genotype-Tissue Expression study (GTEx v6). The second column reports the tissue type based on the relationship to the immune system (main text). The tissues inside each tissue type group are sorted based on number of CDR3 sequences. (3) Histological type of the body site. (4) The median number of 76×2 bp paired-end reads per sample. (5) Number of RNA-Seq samples available via GTEx. Results for (7-40) are presented individually for immunoglobulin heavy chain (IGH), immunoglobulin kappa chain (IGK, immunoglobulin lambda chain (IGK), T cell receptor alpha chain (TCRA), T cell receptor beta chain (TCRB), T cell receptor delta chain (TCRD), and T cell receptor gamma chain (TCRG). (7-8) Median relative abundance of B or T cells within each tissue. (10-16) Median number of distinct CDR3 (clonotypes) is reported per tissue. (18-24) Median number of distinct clonotypes (CDR3) per 1 million RNA-Seq reads (CPM) is reported. (34-40) We used per sample alpha diversity (Shannon entropy) to estimate the diversity of immune repertoire. Median value per tissue is reported.

**Table S3.** The adjusted clonotypic richness of B cells, calculated as the number of distinct amino acid sequences of CDR3 per one million RNA-Seq reads (CPM) normalized by the proportion of the B cell in the sample. We have used SaVant, a transcriptome-based computational method to infer the relative abundance of B cells within each tissue sample based on cell-specific gene signatures (independent of *Ig* transcripts).

| Tissue type | Adjusted CPM (IGH) | Adjusted CPM (IGK) | Adjusted CPM (IGL) |
| --- | --- | --- | --- |
| Adipose - Subcutaneous | 0.31 | 0.10 | 3.33 |
| Adipose - Visceral (Omentum) | 1.22 | 0.18 | 2.92 |
| Adrenal Gland | 0.34 | 0.13 | 3.76 |
| Artery - Aorta | 1.16 | 0.21 | 4.54 |
| Artery - Coronary | 1.19 | 0.21 | 3.86 |
| Artery - Tibial | 0.26 | 0.13 | 8.41 |
| Bladder | 2.47 | 0.23 | 2.86 |
| Brain - Amygdala | 0.10 | 0.08 | 6.78 |
| Brain - Anterior cingulate cortex (BA24) | 0.14 | 0.07 | 7.30 |
| Brain - Caudate (basal ganglia) | 0.13 | 0.07 | 6.58 |
| Brain - Cerebellar Hemisphere | 0.15 | 0.10 | 12.45 |
| Brain - Cerebellum | 0.18 | 0.12 | 13.11 |
| Brain - Cortex | 0.14 | 0.07 | 7.87 |
| Brain - Frontal Cortex (BA9) | 0.13 | 0.08 | 7.89 |
| Brain - Hippocampus | 0.11 | 0.09 | 5.99 |
| Brain - Hypothalamus | 0.10 | 0.07 | 6.27 |
| Brain - Nucleus accumbens (basal ganglia) | 0.12 | 0.07 | 6.86 |
| Brain - Putamen (basal ganglia) | 0.13 | 0.08 | 6.85 |
| Brain - Spinal cord (cervical c-1) | 0.10 | 0.07 | 4.60 |
| Brain - Substantia nigra | 0.09 | 0.07 | 5.49 |
| Breast - Mammary Tissue | 3.91 | 0.37 | 3.40 |
| Cells - EBV-transformed lymphocytes | 0.22 | 0.04 | 1.12 |
| Cells - Transformed | 0.10 | 0.16 | 12.75 |
| fibroblasts |  |  |  |
| Cervix - Ectocervix | 1.75 | 0.34 | 3.18 |
| Cervix - Endocervix | 2.12 | 0.39 | 2.32 |
| Colon - Sigmoid | 0.78 | 0.17 | 5.93 |
| Colon - Transverse | 12.37 | 0.74 | 3.21 |
| Esophagus -<br>Gastroesophageal<br>Junction | 0.16 | 0.11 | 5.62 |
| Esophagus - Mucosa | 2.45 | 0.39 | 6.18 |
| Esophagus -<br>Muscularis | 0.15 | 0.11 | 5.38 |
| Fallopian Tube | 2.79 | 0.33 | 1.86 |
| Heart - Atrial<br>Appendage | 0.26 | 0.13 | 5.78 |
| Heart - Left Ventricle | 0.20 | 0.13 | 6.53 |
| Kidney - Cortex | 1.00 | 0.20 | 3.47 |
| Liver | 1.23 | 0.32 | 5.24 |
| Lung | 3.87 | 0.32 | 2.27 |
| Minor Salivary Gland | 22.36 | 1.18 | 4.37 |
| Muscle - Skeletal | 0.22 | 0.15 | 11.09 |
| Nerve - Tibial | 0.27 | 0.12 | 4.04 |
| Ovary | 0.54 | 0.21 | 6.73 |
| Pancreas | 0.59 | 0.25 | 7.75 |
| Pituitary | 0.27 | 0.13 | 4.44 |
| Prostate | 0.86 | 0.22 | 3.60 |
| Skin - Not Sun<br>Exposed (Suprapubic) | 0.37 | 0.19 | 6.39 |
| Skin - Sun Exposed<br>(Lower leg) | 0.20 | 0.15 | 6.87 |
| Small Intestine -<br>Terminal Ileum | 8.35 | 0.53 | 1.94 |
| Spleen | 10.10 | 0.53 | 1.29 |
| Stomach | 5.95 | 0.50 | 2.93 |
| Testis | 0.13 | 0.08 | 5.45 |
| Thyroid | 1.37 | 0.19 | 4.43 |
| Uterus | 0.52 | 0.23 | 5.33 |

**Table S4.**
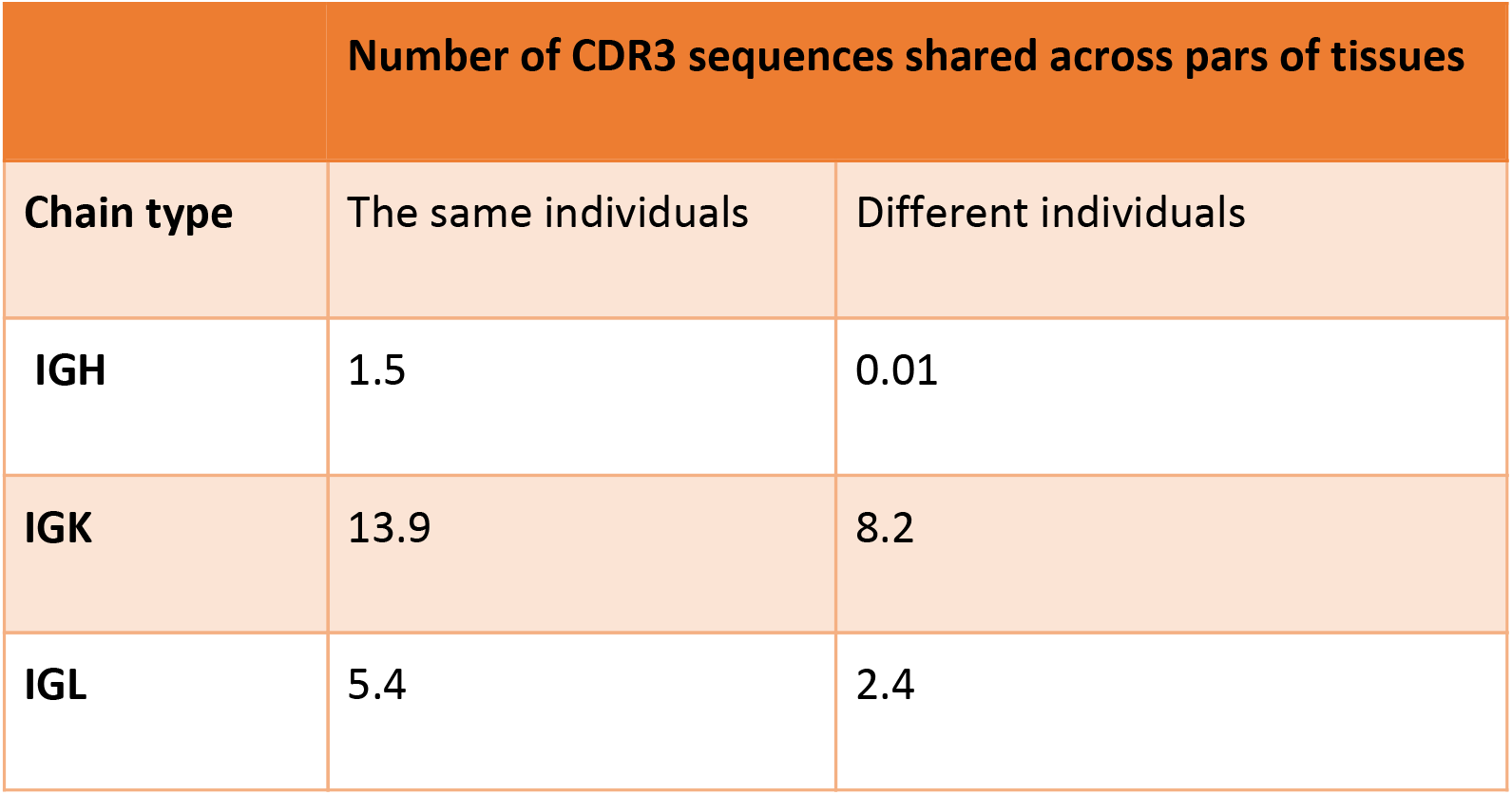
Data used for **Figure 5a** represented as a table. Results are based on pairs of tissues with at least 10 individuals. The number of clonotype sequences shared across pairs of tissues from the same individuals is presented in column 2. The number of clonotype sequences shared across pairs of tissues from different individuals is presented in column 3.

| Number of CDR3 sequences shared across pairs of tissues |  |  |
| --- | --- | --- |
| Chain type | The same individuals | Different individuals |
| IGH | 1.5 | 0.01 |
| IGK | 13.9 | 8.2 |
| IGL | 5.4 | 2.4 |

**Table S5.** The top 25 genes with the highest proportional median value for CD19+ B cells, CD4+ T cells, and CD8+ T cells.

| B cell signature |  | CD4+ T cell Signature |  | CD8+ T cell Signature |  |
| --- | --- | --- | --- | --- | --- |
| **HLA-DQA1** | major histocompatibility complex, class II, DQ alpha 1 | **CD3D** | CD3d molecule, delta (CD3-TCR complex) | **LCK** | lymphocyte-specific protein tyrosine kinase |
| **HLA-DQA2** | major histocompatibility complex, class II, DQ alpha 2 | **PTPRC** | protein tyrosine phosphatase, receptor type, C | **CD247** | CD247 molecule |
| **HLA-DMA** | major histocompatibility complex, class II, DM alpha | **LCK** | lymphocyte-specific protein tyrosine kinase | **CD3D** | CD3d molecule, delta (CD3-TCR complex) |
| **HLA-DOB** | major histocompatibility complex, class II, DO beta | **CD247** | CD247 molecule | **PIK3CD** | phosphoinositide-3-kinase, catalytic, delta polypeptide |
| **CXCR4** | chemokine (C-X-C motif) receptor 4 | **PIK3CD** | phosphoinositide-3-kinase, catalytic, delta polypeptide | **CXCR4** | chemokine (C-X-C motif) receptor 4 |
| **SELL** | selectin L | **CXCR4** | chemokine (C-X-C motif) receptor 4 | **GZMA** | granzyme A (granzyme 1, cytotoxic T-lymphocyte-associated serine esterase 3) |
| **CD79A** | CD79a molecule, immunoglobulin-associated alpha | **IL7R** | interleukin 7 receptor | **CD48** | CD48 molecule |
| **ISG20** | interferon stimulated exonuclease gene 20kDa | **SELL** | selectin L | **IL7R** | interleukin 7 receptor |
| **MS4A1** | membrane-spanning 4-domains, subfamily A, member 1 | **ICAM3** | intercellular adhesion molecule 3 | **SELL** | selectin L |
| **CD37** | CD37 molecule | **IL10RA** | interleukin 10 receptor, alpha | **LTB** | lymphotoxin beta (TNF superfamily, member 3) |
| **CD48** | CD48 molecule | **CCR7** | chemokine (C-C motif) receptor 7 | **CTSW** | cathepsin W |
| **LTB** | lymphotoxin beta (TNF superfamily, member 3) | **LTB** | lymphotoxin beta (TNF superfamily, member 3) | **HMHA1** | histocompatibility (minor) HA-1 |
| **P2RX5** | purinergic receptor P2X, ligand-gated ion channel, 5 | **CD48** | CD48 molecule | **CORO1A** | coronin, actin binding protein, 1A |
| **LAPTM5** | lysosomal protein transmembrane 5 | **LAPTM5** | lysosomal protein transmembrane 5 | **KLRB1** | killer cell lectin-like receptor subfamily B, member 1 |
| **PLAC8** | placenta-specific 8 | **HMHA1** | histocompatibility (minor) HA-1 | **NKG7** | natural killer cell group 7 sequence |
| **POU2AF1** | POU class 2 associating factor 1 | **CORO1A** | coronin, actin binding protein, 1A | **PLAC8** | placenta-specific 8 |
| **TCL1A** | T-cell leukemia/lymphoma 1A | **KLRB1** | killer cell lectin-like receptor subfamily B, member 1 | **DENND2D** | DENN/MADD domain containing 2D |
| **FAIM3** | Fas apoptotic inhibitory molecule 3 | **PLAC8** | placenta-specific 8 | **FAIM3** | Fas apoptotic inhibitory molecule 3 |
| **AL928768.3** | lincRNA | **DENND2D** | DENN/MADD domain containing 2D | **KLRC4** | killer cell lectin-like receptor subfamily C, member 4 |
|  |  | **FAIM3** | Fas apoptotic inhibitory molecule 3 |  |  |
|  |  | **GMFG** | glia maturation factor, gamma |  |  |

